# The evolutionary interplay between self-fertilisation and sex allocation in flowering plants

**DOI:** 10.64898/2026.10.01.755674

**Authors:** Thomas Lesaffre, John R. Pannell, Charles Mullon

## Abstract

Flowering plants vary in sex allocation (resource division between male and female function) and in selfing rate. These traits should evolve together: selfing changes fitness returns from pollen and ovules, whereas sex allocation changes the pollen available for selfing and export. Yet most models allow only one trait to evolve, leaving their evolutionary feedback unresolved. Here we model the joint evolution of sex allocation, pollen export and maternal control of competition between self- and cross-pollen. We show that selfing and sex allocation strongly influence one another, as selection favours associations between sex allocation and mating-system traits. When maternal control is fixed to recover classical mass-action, evolution can lead to the stable coexistence of selfing hermaphrodites and males specialised in pollen export. This androdioecy persists because hermaphrodites cannot prevent fertilisation by males. When maternal control of pollen competition also evolves, we find that where a mixed-mating equilibrium exists, selection again favours the emergence of androdioecy. As hermaphrodites evolve to favour their own pollen, however, they progressively exclude pollen from males, driving them extinct. The population may then return to mixed mating and repeat the process, or evolve complete selfing. We discuss these results with reference to known variation in the mating and sexual systems of plants.

## 1 Introduction

Flowering plants vary widely in their sexual system, that is in how male and female functions are distributed across individuals (Renner and Ricklefs, 1995; Renner, 2014; Käfer et al., 2017). Most species are hermaphroditic, meaning that each individual produces both pollen and ovules, but the resources allocated to each function can still vary substantially (Cruden, 1977; Cruden and Lyon, 1985; Brunet, 1992; Jürgens et al., 2002). Other species have evolved polymorphic sexual systems including dioecy, in which individuals are either male or female, as well as rarer systems in which hermaphroditic and unisexual individuals (males or females) coexist in the same population (Renner, 2014; Käfer et al., 2017). This diversity reflects the range of selective pressures acting on plant sex allocation, i.e., how plants divide their reproductive resources between male and female function (Freeman et al., 1997; Käfer et al., 2017; Pannell and Jordan, 2022).

The evolution of plant sex allocation depends on ecological features that influence the fitness returns from producing pollen and ovules (Charnov et al., 1976; Charnov, 1982). These features for example include the density of potential mates and competitors through each function, plant size, and interactions with animal pollinators, seed dispersers and predators (for reviews, see Bawa, 1980; Charlesworth, 1993, 1999; Freeman et al., 1997; De Jong and Klinkhamer, 2005; Barrett and Harder, 2017; Leonard, 2018; Pannell and Jordan, 2022). Among them, the population’s mating system – the dynamics of pollen exchange between flowers that determine who fertilises whom (Barrett and Harder, 2017) – is especially influential.

Self-fertilisation (selfing hereafter), a common mating strategy in hermaphrodites whereby individuals fertilise their own ovules, is thought to have profound consequences for sex allocation evolution. First, selfing changes expected fitness returns through male and female function in a way that favours female-biased sex allocation (Charlesworth and Charlesworth, 1981; Zhang, 2000; Lesaffre et al., 2025). This is because selfing simultaneously : (i) decreases expected fitness gains as a pollen donor by reducing the number of ovules available for outcrossing; and (ii) increases expected fitness returns through seeds when inbreeding depression – the decrease in relative fitness of inbred progeny owing to homozygosity for recessive deleterious mutations (Charlesworth and Charlesworth, 1987) – is weak relative to the trans-mission advantage that selfing entails (a selfing parent acts as both mother and father to its offspring, and so transmits twice as many gene copies per seed on average; Fisher, 1941). Second, selfing is thought to play a central role in many evolutionary transitions from combined to separate sexes that have occurred in flowering plants (Charlesworth, 1999; De Jong and Geritz, 2001; Pannell and Jordan, 2022). Models show that selfing and inbreeding depression can facilitate the evolution of separate sexes, either through the step-wise invasion of hermaphroditic populations by sterility mutations creating unisexual females and males (Charlesworth and Charlesworth, 1978), or via the gradual emergence of specialised male and female morphs (Lesaffre et al., 2024).

Existing theory has clarified how selfing affects sex allocation evolution, but most of it treats the mating system as fixed, despite the fact that selfing rates are known to evolve rapidly (Takebayashi and Morell, 2001; Goldberg et al., 2010; Barrett and Harder, 2017). The mechanisms shaping selfing evolution have been extensively discussed (Goodwillie et al., 2005; Clo et al., 2025). Early theoretical treatments show that the opposition between the transmission advantage (Fisher, 1941) and inbreeding depression generates a threshold effect: selection favours either complete outcrossing or complete selfing depending on the severity of inbreeding depression, so that mixed mating, where individuals self-fertilise at an intermediate rate, should not be stable (Lande and Schemske, 1985). This insight came from models treating inbreeding depression as a parameter, but studies that explicitly modelled the genetic basis of inbreeding depression reached similar conclusions (e.g., Charlesworth et al., 1990, 1991; Porcher et al., 2009; Lande and Porcher, 2015; Abu Awad and Billiard, 2017; Abu Awad and Roze, 2020). Additional mechanisms may therefore be needed to explain the widespread occurrence of mixed mating in flowering plants (Goodwillie et al., 2005; Whitehead et al., 2018).

Mixed mating may often simply result from accidental fertilisations within and between flowers on the same plant owing to imperfect outcrossing mechanisms (de Jong et al., 1993; Barrett, 2003), but it may also be favoured by selection through several ecological mechanisms (see Goodwillie et al., 2005; Clo et al., 2025 for reviews). These include density- or environment-dependent variation in inbreeding depression (Cheptou and Mathias, 2001; Cheptou and Dieckmann, 2002; Cheptou and Donohue, 2011), limited dispersal of pollen and seed (Ronfort and Couvet, 1995) and uncertain pollination (Morgan and Wilson, 2005). It is however pollen discounting – the reduction in male mating success through outcrossing that can be caused by traits increasing the selfing rate (Harder and Wilson, 1998) – that has received the most attention. Pollen discounting has been modelled in two ways depending on the mode of selfing considered. In models of prior selfing, where selfing occurs before any opportunity for outcrossing (Lloyd, 1979), pollen discounting is introduced by making individuals’ contribution to the outcross pollen pool a function of their selfing rate (e.g., Nagylaki, 1976; Johnston, 1998; Abu Awad and Roze, 2020), reflecting a form of sexual interference whereby floral traits that increase selfing compromise pollen export (Barrett, 2002). By contrast, pollen discounting occurs mechanistically in mass-action models of competing selfing (Lloyd, 1979), where self- and cross-pollen compete for fertilisation. In these models, the selfing rate is determined by the proportion of self-pollen among all pollen deposited on the stigma. Pollen remaining on the plant for selfing is necessarily removed from the contribution individuals make to the outcross pollen pool, resulting in pollen discounting (Holsinger et al., 1984; Holsinger, 1991; Porcher and Lande, 2005). In both prior and competing selfing models, pollen discounting favours intermediate selfing rates because it generates a cost to self-fertilisation that increases non-linearly with the selfing rate (Holsinger, 1991; Johnston, 1998; Porcher and Lande, 2005).

Pollen discounting is therefore thought to be important for selfing evolution, but most of the theory on which this is based assumes fixed sex allocation. All individuals are thus assumed to produce the same number of pollen grains and ovules and so to experience pollen discounting similarly. If sex allocation also evolved, however, the extent of pollen discounting could depend on how much an individual allocates to pollen production. Individuals allocating more heavily to pollen may need to spend a smaller proportion of their pollen to achieve a given selfing rate, so that the relative decrease in their male outcrossing success with the selfing rate (i.e., pollen discounting) might be less severe. Furthermore, empirical evidence suggests that individual selfing rates in fact often vary with individual sex allocation (Damgaard and Abbott, 1995; Chen and Pannell, 2024, 2025).

Selfing and sex allocation might therefore often evolve jointly, yet this joint evolution has rarely been considered. To date, the only model addressing this question is that of Zhang (2000), who studied the joint evolution of sex allocation and pollen export under a mass-action model of competing selfing, where self- and cross-pollen arrive simultaneously on the stigma and compete for fertilisation (Lloyd, 1979; Holsinger, 1991). Zhang (2000) found that although partial selfing favours female-biased sex allocations (as expected, Charlesworth and Charlesworth, 1981), the equilibrium rate of pollen export favoured by selection is independent of sex allocation, suggesting that the mating system influences sex allocation evolution but is not influenced in return. However, the analyses of Zhang (2000) are limited to directional selection and therefore do not address what happens once the population has converged to the equilibrium. In particular, they do not determine whether this equilibrium remains stable or gives rise to evolutionary branching, whereby distinct types of individuals emerge and become increasingly differentiated, potentially leading to a polymorphic sexual system (Geritz et al., 1998; Pásztor et al., 2016; Avila and Mullon, 2023). Moreover, Zhang (2000)’s model relies on the mass-action assumption, which constrains selfing to be strictly competing and so excludes other modes of self-fertilisation such as prior selfing, where self-pollen fertilises ovules before any opportunity for outcrossing; or delayed selfing, where self-fertilisation only occurs after outcrossing has failed (Lloyd, 1979).

Here, we model the joint evolution of sex allocation and the mating system, allowing for different modes of selfing and investigating what happens beyond directional selection. We introduce an extension of the mass-action model where maternal plants express a trait that allows them to manipulate pollen competition either in favour of or against their own pollen, so that the model encompasses the three modes of selfing identified by Lloyd (1979): prior, competing and delayed. First keeping this trait fixed so that we recover the classical mass-action model (Zhang, 2000), we show that the joint evolution of sex allocation and pollen export can in fact lead to a form of androdioecy where males and partially selfing hermaphrodites coexist. We then let the pollen-competition trait evolve jointly with sex allocation and pollen export, and find that this leads to recurrent episodes of sexual specialisation followed by its collapse, ultimately driving the population to complete selfing through transient androdioecy. Altogether, our results shine light on potential evolutionary feedbacks between plant mating and sexual systems that had so far been overlooked.

## 2 Model and analytical approach

### 2.1 Biological scenario

We consider a population composed of a large and constant number *N* of hermaphroditic plants with an annual life-history, so that generations are non-overlapping. We assume throughout our analyses that plants are haploid with a reduced diploid phase after gamete fusion: ovule fertilisation results in the production of diploid zygotes that rapidly undergo meiosis to give rise to two haploid seeds each. This assumption simplifies the analyses. It allows us to track the evolution of phenotypes in the absence of genetic constraints on the trait combinations that evolve, with in principle no impact on adaptation in the long-term (e.g., Geritz and Kisdi, 2000; Lesaffre et al., 2025; Lehmann and Mullon, 2026). We address the influence of this simplified genetic architecture on our results in the Discussion.

Each individual expresses three continuous traits **z** = (*z*_1_, *z*_2_, *z*_3_), each varying between zero and one. The first trait *z*_1_ is sex allocation, defined as the proportion of its reproductive resources that an individual devotes to its female function. The second trait *z*_2_ is the pollen export rate, that is the proportion of its pollen that an individual disperses to fertilise ovules on other plants (while the rest remains on the plant for selfing, as assumed under the mass-action model; Holsinger, 1991). The third trait *z*_3_, which we refer to as the self-pollen advantage hereafter, allows maternal plants to manipulate the outcome of pollen competition either in favour of or against their own pollen (this will be made explicit in eq. (2a) below). These three traits determine individuals’ reproductive behaviour as they pass through the following life cycle events.

#### (i) Sexual development

Plants first grow and mature sexually. Plant *i* ∈ {1, …, *N*} allocates a fraction *z*_1,*i*_ ∈ [0, 1] of its resources to its female function, and the remaining fraction 1 − *z*_1,*i*_ to its male function, resulting in the production of

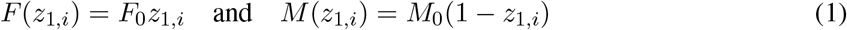

ovules and pollen grains, respectively. Throughout, we assume that the number of ovules is large relative to the number of breeding spots (*N* ≪ *F*_0_), and that there is always enough pollen for all ovules to be fertilised (*F*_0_ ≪ *M*_0_), i.e., no pollen limitation.

#### (ii) Mating

Following pollen and ovule production, individuals export a fraction *z*_2_ ∈ [0, 1] of their pollen to fertilise ovules on other plants, and the rest (1−*z*_2_) remains on the plant for self-fertilisation (Holsinger, 1991; Zhang, 2000; Porcher and Lande, 2005). A fraction *c* ∈ [0, 1] of the exported pollen is lost during dispersal, and the successful fraction 1−*c* is distributed equally among the *N* −1 other plants. The pollen export rate *z*_2_ thus determines the absolute amount of pollen transferred within and between plants. In animal-pollinated species, this rate could be influenced by traits that modulate pollinator attraction (e.g., flower colour or odour), as higher visitation rates should result in higher rates of pollen removal (Johnson and Harder, 2023). In wind-pollinated plants, the pollen export rate might be affected by morphological traits that affect the removal of pollen from anthers by gusts of wind, such as peduncle length or features of pollen morphology (Niklas, 1985).

The self- and cross-pollen received by each plant compete for ovule fertilisation, and this competition is biased by the maternal plant either in favour of or against its own pollen according to its self-pollen advantage *z*_3,*i*_. We scale this effect such that competition is unbiased when *z*_3,*i*_ = 1*/*2, so that pollen grains from all sources are equally competitive (as envisioned under mass-action), whereas *z*_3,*i*_ *>* 1*/*2 and *z*_3,*i*_ *<* 1*/*2 favour and oppose self-fertilisation, respectively. The self-pollen advantage *z*_3_ thus modulates the relative competitiveness of self- and cross-pollen grains for fertilisation. This trait could be influenced by features of floral morphology that affect the distance between anthers and pistils (herkogamy), or by aspects of floral phenology that affect the timing of anther dehiscence relative to the opening of flowers (dichogamy), both of which may evolve to give either a head-start or a handicap to self-pollen in the race to ovules (Karron et al., 1997; Goodwillie and Weber, 2018). In addition, it could reflect less conspicuous mate choice mechanisms that occur in the style, as maternal plants can hinder (or boost) the progression of pollen tubes from particular sources relative to others in the race towards ovules (Tonnabel et al., 2021; Chenin et al., 2026a,b), as is the case in many self-incompatibility systems for example (Takayama and Akira, 2005; Durand et al., 2020).

Overall, the selfing rate *α* (**z**_*i*_) of individual *i* expressing phenotype **z**_*i*_ = (*z*_1,*i*_, *z*_2,*i*_, *z*_3,*i*_) when the *N* individuals in the population express phenotypes **z**_1_, **z**_2_, …, **z**_*N*_ is given by

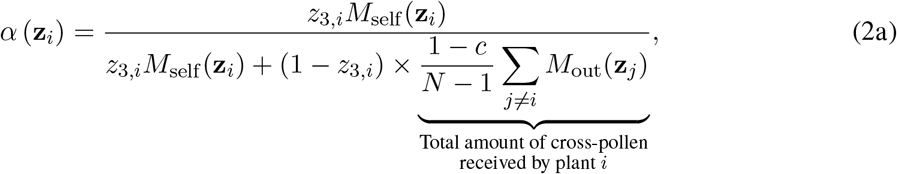

and the probability that individual *i* fertilises an ovule of individual *j*≠ *i* is

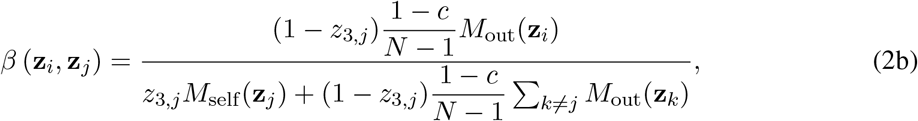

where

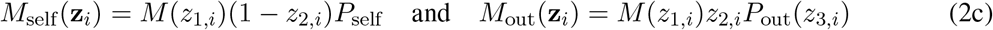

respectively give the amount of self-pollen transferred to plant *i*’s own stigmas and the amount of pollen exported away by plant *i*. In eq. (2c), *P*_self_ ∈ [0, 1] is a parameter that captures the efficiency of within-plant pollen transfers, while *P*_out_(*z*_3,*i*_) ∈ [0, 1] captures the efficiency of pollen export, which depends on the self-pollen advantage *z*_3,*i*_ of the individual.

We assume that *P*_out_(*z*_3,*i*_) decreases with *z*_3,*i*_, reflecting the notion that expressing floral features that favour self-pollination might decrease pollen export efficiency, and so lead to pollen discounting due to a form of sexual interference (Barrett, 2002). Mechanistically, this could result from changes to floral morphology that cause self-pollen to be deposited earlier on pistils, and in doing so decrease the efficiency with which pollen is released from anthers and/or exported away from the plant. For example, reduced herkogamy may lead self-pollen to reach the stigma more rapidly, but it may cause the pistil to get in the way of efficient pollen release (Karron et al., 1997; Opedal, 2018). Such a morphological change would give the deposited self-pollen a head-start in the race to ovules, but reduce the amount exported, resulting in pollen discounting.

Since *M*_self_ (**z**_*i*_) and *M*_out_(**z**_*i*_) appear exclusively in ratios in eqs. (2a) and (2b), only their relative scaling matters. Without loss of generality, we may therefore divide both by *P*_self_ so that they become

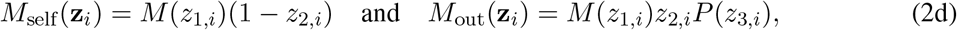

where *P* (*z*_3,*i*_) = *P*_out_(*z*_3,*i*_)*/P*_self_ is the relative efficiency of pollen export, which we will work with from now on.

#### (3) Seed maturation

Fertilisation results in the production of diploid zygotes. Outcrossed zygotes become viable with probability one, meanwhile self-fertilised zygotes suffer from inbreeding depression, becoming viable with probability 1 − δ, where δ ∈ [0, 1] is a parameter measuring the severity of inbreeding depression. Following many authors (e.g., Lande and Schemske, 1985; Lloyd, 1992; Morgan et al., 1997; Rausher and Chang, 1999; Zhang, 2000; Cheptou and Mathias, 2001; Cheptou and Dieckmann, 2002; Tsitrone et al., 2003; Morgan and Wilson, 2005, see Goodwillie et al., 2005 and Clo et al., 2025 for reviews), we therefore assume inbreeding depression to be fixed. We discuss the influence of relaxing this assumption on our results at the end of the paper. Viable diploid zygotes immediately undergo meiosis to give rise to two haploid seeds each.

#### (4) Germination and recruitment

All adults die following seed maturation. The produced seeds then germinate and compete to fill the *N* breeding spots in the population.

We assume that sex allocation *z*_1_, the pollen export rate *z*_2_ and the self-pollen advantage *z*_3_ are encoded by a single pleiotropic locus at which alleles may encode any trait combination **z** = (*z*_1_, *z*_2_, *z*_3_) ∈ [0, 1]^3^ following a continuum-of-alleles model (Kimura, 1965; Walsh and Lynch, 2018). Mutations occur at a small fixed rate at this locus, adding small, unbiased and independent deviations onto the trait values encoded by the mutated copy.

### 2.2 Invasion analysis

When mutations are rare and have small effects on phenotype, evolution proceeds gradually and its trajectory can be inferred from an invasion analysis (Geritz et al., 1998; Pásztor et al., 2016; Avila and Mullon, 2023). The basis to such analysis is the invasion fitness *W* (**z**_*•*_, **z**) of a mutant **z**_*•*_ = (*z*_1*•*_, *z*_2*•*_, *z*_3*•*_) ap-pearing as a single copy in a resident population otherwise fixed for **z** = (*z*_1_, *z*_2_, *z*_3_). In a homogeneous population, *W* (**z**_*•*_, **z**) is given by the number of successfully recruited mutant gene copies produced by a rare focal mutant over a full iteration of the life cycle. In the context of our model, we show in Appendix A.1.1 that invasion fitness can be expressed as

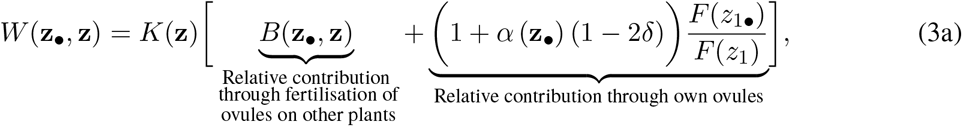

where *K*(**z**) *>* 0 is a positive factor that captures density regulation, and

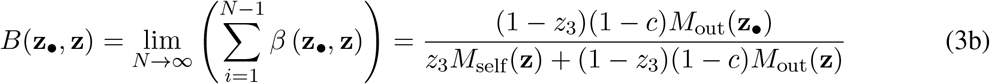

measures the relative contribution that the mutant makes to the next generation by fertilising ovules on other plants in a large population, i.e., the mutant’s total siring success as a pollen donor (see eq. A1 in Appendix A.1.1 for more details).

The term between square brackets in eq. (3a) measures the relative number of mutant copies contributed by the mutant to the next generation, which is composed of two parts. The first part, *B*(**z**_*•*_, **z**), measures the contribution it makes as a pollen donor (eq. 3b). The second part corresponds to the relative contribution the mutant makes through its own ovules, and can be understood as follows. The mutant produces *F* (*z*_1*•*_) mutant ovules, a proportion *α* (**z**_*•*_) of which get self-fertilised. The resulting self-fertilised zygotes become viable with probability 1 − δ, and divide into two haploid mutant seeds when they do so, yielding 2(1 − δ) mutant copies per ovule on average. The remaining ovules 1 − *α* (**z**_*•*_) get fertilised by resident pollen, become viable with probability 1 and then divide to give rise to a resident and a mutant seed each (this reduced diploid phase therefore captures the two-fold transmission advantage of selfing, Fisher, 1941). As a result, the average number of mutant copies transmitted per ovule by the mutant through its female function is given by 2(1−δ)*×α* (**z**_*•*_)+1*×*(1−*α* (**z**_*•*_)) = 1+*α* (**z**_*•*_) (1−2δ), and the total number of mutant copies contributed by the mutant to the offspring pool is equal to *F* (*z*_1*•*_)*×* (1+*α* (**z**_*•*_) (1−2δ), which is then scaled by resident female fecundity *F* (*z*_1_).

#### Directional selection

Equation (3) can be used to infer the joint evolutionary dynamics of any combination of the three traits, which occur in two phases (Geritz et al., 1998; Leimar, 2009). At first, an initially monomorphic population experiences directional selection, which causes positively selected mutations to rapidly sweep to fixation, so that the population travels through phenotypic space in the direction favoured by selection while remaining largely monomorphic. The direction of selection on a trait *z*_*v*_, *v* ∈ {1, …, *V*} (where *V* denotes the number of jointly evolving traits), is given by the selection gradient, which one can obtain from invasion fitness (eq. 3) as

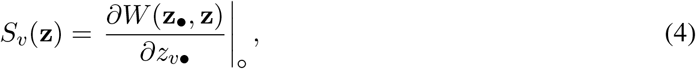

where the |_*○*_ notation indicates that the derivative is evaluated at the resident strategy (**z**_*•*_ = **z**). A positive selection gradient indicates that directional selection favours an increase in the trait, whereas a negative gradient indicates that it favours a decrease. Through the recurrent fixation of mutations taking the population in the direction favoured by selection, the population may eventually attain a singular strategy 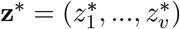 at which directional selection vanishes, i.e.,

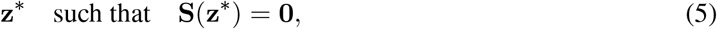

where vector **S**(**z**) = (*S*_*v*_(**z**))_1⩽*v*⩽*V*_ collects the selection gradients on each evolving trait. The mathematical conditions for convergence to a singular strategy are treated in Leimar (2009) (see also Appendix A.1.2.1 here).

#### Disruptive selection

Once the population has converged to a singular strategy, selection may either be stabilising around **z**^∗^, in which case the population remains monomorphic for these trait values and evolutionary dynamics grind to a halt; or selection may be disruptive, potentially leading to evolutionary branching and the gradual emergence of polymorphism (Geritz et al., 1998; Pásztor et al., 2016; Avila and Mullon, 2023). Whether selection is stabilising or disruptive around **z**^∗^ is assessed from the Hessian matrix **H** (**z**^∗^), which is a symmetrical *V × V* matrix the (*u, v*)-entry of which is given by

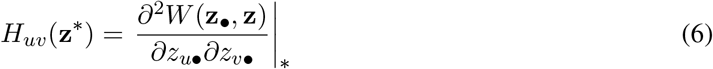

for *u, v* ∈ {1, …, *V*}, where the |_∗_ notation indicates that the derivative is evaluated at the singular strategy (**z**_*•*_ = **z** = **z**^∗^). Disruptive selection requires the leading eigenvalue of **H** (**z**^∗^) to be positive (Leimar, 2009; Débarre et al., 2014; Geritz et al., 2016).

The entries of the Hessian matrix **H** (**z**^∗^) carry information regarding selection on each trait and their combination. Each diagonal entry (*u* = *v* in eq. 6) gives the quadratic selection coefficients on each trait, informing us on the nature of selection on the trait taken independently (Lande and Arnold, 1983). Specifically, a positive quadratic coefficient *H*_*vv*_(**z**^∗^) *>* 0, *v* ∈ {1, …, *V*}, indicates that selection is disruptive on trait *z*_*v*_, and a negative coefficient indicates that it is stabilising. The off-diagonal entries (*u*≠ *v* in eq. 6), meanwhile, give the correlational selection coefficient between pairs of traits. These tell us whether selection favours a positive (*H*_*uv*_(**z**^∗^) *>* 0) or negative (*H*_*uv*_(**z**^∗^) *<* 0) correlation between two traits within individuals when they evolve jointly (Lande and Arnold, 1983).

A more detailed description of the method is given in Appendix A.1 and an in-depth analysis of the model is available in Appendices A.2 and A.3. We summarise our key findings below.

## 3 The joint evolution of sex allocation and pollen export enables sexual specialisation

As a starting point, we assume that the self-pollen advantage is fixed at *z*_3_ = 1*/*2, so that pollen competition is unbiased, and that there is no sexual interference (*P* (*z*_3_) = 1). This recovers a classical mass-action model (Holsinger, 1991). In this setting, we analyse the joint evolution of sex allocation *z*_1_ and pollen export *z*_2_, as previously studied by Zhang (2000), extending their results to consider whether polymorphism can emerge.

### 3.1 Directional selection leads to outcrossing or mixed mating depending on inbreeding depression

Using eq. (4), we show in Appendix A.2.1 that the selection gradients on sex allocation *z*_1_ and on pollen export *z*_2_ simplify to

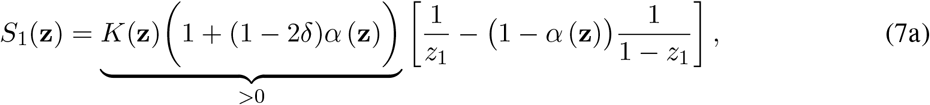

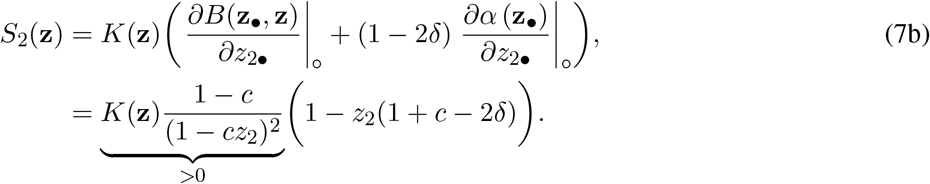

Analysing these gradients jointly reveals that directional selection leads to one of two evolutionary equilibria depending on the severity of inbreeding depression (δ) relative to the costs of pollen dispersal (*c*), with (*z*_1_, *z*_2_) converging either to

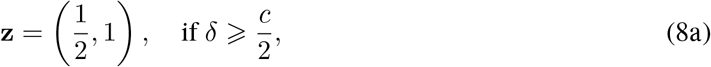

or to the singular strategy

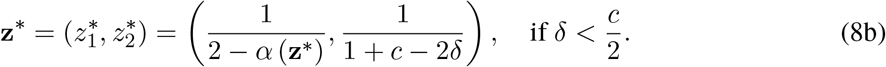

The first equilibrium corresponds to a population of completely outcrossing hermaphrodites, whereas the second consists of partially selfing, female-biased hermaphrodites, where the implicit expression for *z*_1_^∗^ simplifies to 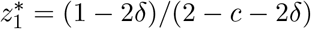.

The reason why these two outcomes arise is apparent from the selection gradients. The gradient on sex allocation (eq. 7a) always vanishes at 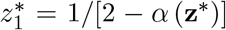, showing that the equilibrium sex allocation is entirely determined by the selfing rate. Under complete outcrossing *α* (**z**^∗^) = 0, selection favours equal investment into male and female function, as predicted by classical theory (Charnov et al., 1976; Charnov, 1982). When selfing occurs (*α* (**z**^∗^) *>* 0), the equilibrium becomes female-biased because incomplete pollen dispersal reduces the opportunities for fitness gains through pollen export (see eq. A20a and associated discussion in Appendix A.2.1 for more details). Meanwhile, the selection gradient on pollen export (eq. 7b) is independent of sex allocation *z*_1_, consistent with previous findings (Zhang, 2000). It reveals a threshold determined by the balance between inbreeding depression δ and dispersal cost *c*.

When δ ⩾ *c/*2, selection favours complete pollen export. When δ *< c/*2, selection instead favours the intermediate pollen export rate 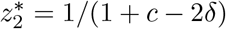, which decreases as pollen dispersal becomes more costly and increases as inbreeding depression becomes stronger. Why selection favours an intermediate 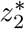 and so mixed mating can be understood by considering a fully selfing population, in which individuals export no pollen (*z*_2_ = 0). In such a population, a rare mutant exporting some pollen will maintain the same selfing because it is not receiving any pollen from others, therefore maintaining the same fitness gain as others through its own ovule; while simultaneously gaining reproductive success by fertilising ovules on other plants (who cannot avoid it under strict mass-action). It will thus always be at a fitness advantage.

These results recover those of Zhang (2000) concerning directional selection. In particular, the selection gradient on pollen export is independent of sex allocation. We next investigate whether this independence persists at the level of disruptive selection, or if the joint evolution of sex allocation and pollen export can in fact favour the emergence of polymorphic sexual systems.

### 3.2 Emergence of androdioecy through evolutionary branching

When δ ⩾ *c/*2, eq. (7) shows that pollen export remains under directional selection at the boundary of trait space (i.e., *S*_2_(1*/*2, 1) *>* 0). The model thus reduces to an outcrossing hermaphroditic population with linear fitness returns through each sexual function. In this situation, equal investment into male and female function,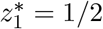 is under stabilising selection (i.e., *H*_11_(1*/*2, 1) ⩽ 0) and therefore constitutes an evolutionarily stable strategy, as shown previously (Charnov, 1982; Lesaffre et al., 2024, 2025).

When δ *< c/*2, directional selection leads the population to the mixed mating singular strategy given in eq. (8b), where selection is now determined by the Hessian matrix **H** (**z**^∗^) (eq. 6), which here takes the form of a symmetric 2 *×* 2 matrix since only two traits are evolving, with entries

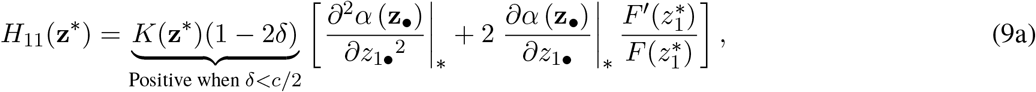

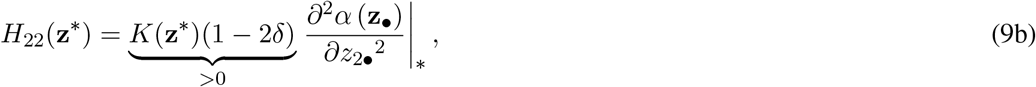

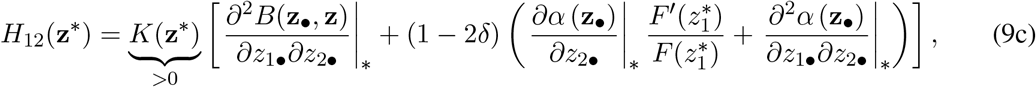

where *F*′(*z*_1_) = d*F* (*z*_1_)*/*d*z*_1_ denotes the first derivative of female fecundity.

The diagonal elements *H*_11_(**z**^∗^) and *H*_22_(**z**^∗^) are both negative, indicating that quadratic selection on sex allocation and on pollen export is stabilising when either trait evolves in isolation from the other. For sex allocation (eq. 9a), this is due to two effects. The first is given by the second derivative of the selfing rate (*α*) with respect to mutant sex allocation, which is negative because the individual’s own pollen increasingly competes with itself for self-fertilisation as the amount produced increases, leading to diminishing returns on the selfing rate (analogous to a form of local sperm/mate competition, Hamilton, 1967; Schärer, 2009). The second effect captures the fitness consequences of change in sex allocation through its simultaneous effects on female fecundity (*F*) and fitness returns per ovule, as influenced by the selfing rate (*α*). Since any change in sex allocation has opposite effects on these two quantities, this effect is also negative. For pollen export (eq. 9b), the sign of quadratic selection is determined by the second derivative of the selfing rate with respect to pollen export, which is also negative. As the proportion of pollen exported decreases, competition amongst the focal individual’s own pollen grains intensifies, again leading to diminishing returns on the selfing rate.

When sex allocation and pollen export evolve jointly, their evolutionary trajectory also depends on correlational selection, which is given by the off-diagonal element *H*_12_(**z**^∗^) (eq. 9c). Overall, we find that this element is always negative, indicating that selection favours a negative correlation between the two traits. This result arises because both components of *H*_12_(**z**^∗^) favour such a correlation. The first component captures the effect of a joint increase of the two traits on success as a pollen donor. It is always negative because increasing allocation to female function *z*_1_ reduces siring success a pollen donor, and the effect of this reduction on fitness is amplified as the focal individual’s pollen export increases (intuitively, individuals that gain more fitness through pollen dispersal suffer a bigger fitness cost from reducing pollen production). The second component, which is proportional to 1 − 2δ, captures the interaction between *z*_1_ and *z*_2_ on fitness gains through the focal individual’s own ovules. Its first part captures the joint effect of a change in pollen export on the selfing rate and a change in sex allocation on relative ovule production. Because increasing pollen export has a negative effect on the selfing rate and increasing sex allocation has a positive effect on ovule production, this part is negative and therefore also favours a negative correlation between pollen export and sex allocation. The second part is the cross-derivative of the selfing rate with respect to both traits. Although it may be either positive or negative depending on parameter values, we find that it is always quantitatively small relative to the other components of eq. (9c) (see Appendix A.2.2 for a detailed discussion). Selection therefore always favours individuals allocating more to their female function to export less pollen, and individuals favouring their male function to export more.

The quadratic selection coefficients in eqs. (9a) and (9b) show that if either trait evolved in isolation from the other, the population would remain monomorphic under stabilising selection. When the two traits coevolve, however, correlational selection can generate disruptive selection if it is strong relative to quadratic selection on both traits, specifically if *H*_12_(**z**^∗^)^2^ *> H*_11_(**z**^∗^)*H*_22_(**z**^∗^). We show in Appendix A.2.2 that this condition is always satisfied, indicating that the Hessian matrix always has a positive eigenvalue (Fig. 1A) and selection favours the emergence of polymorphism through evolutionary branching. Intuitively, this result can be understood by keeping in mind that inbreeding depression is weak here (δ *< c/*2), so that self-fertilised seeds provide greater average fitness returns than outcrossed seeds. In this setting, an individual allocating more resources to ovule production is also able to increase the fitness returns it gains per ovule by simultaneously reducing its pollen export rate. This creates a multiplicative increase in fitness gains through female function that favours specialisation into the production of self-fertilised seeds. However, as individuals cannot avoid fertilisation by others under strict mass-action, the emergence of such a female-biased type also creates more opportunities for fitness gains as a pollen donor, favouring individuals that simultaneously allocate more resources to their male function and export pollen at a higher rate.

**Figure 1:**
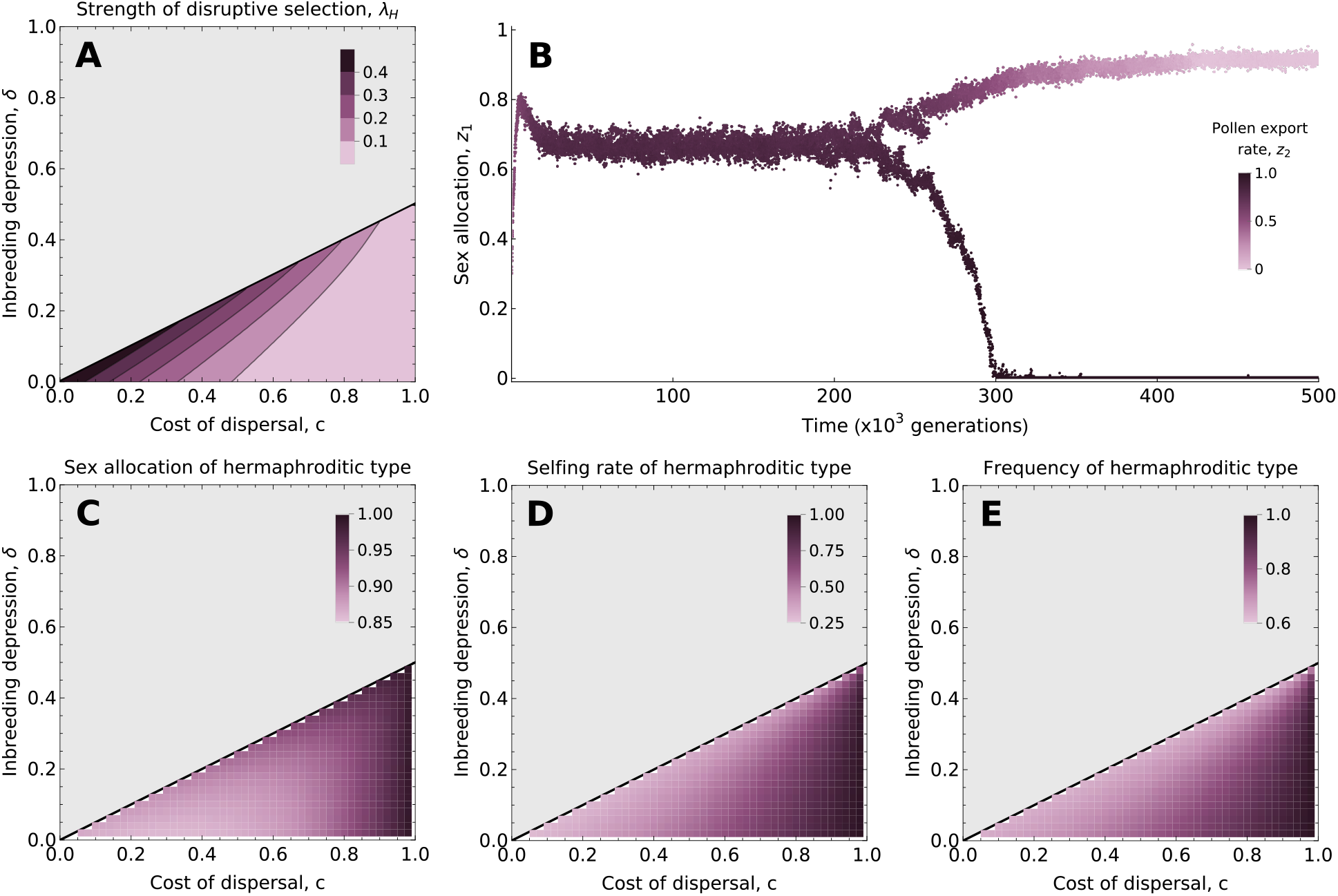
Joint evolution of sex allocation and pollen export produces stable androdioecy. **A** Strength of disruptive selection as given by the leading eigenvalue of the Hessian matrix as a function of the cost of pollen dispersal (*c*) and inbreeding depression (δ). Values are only represented where δ *< c/*2, as the population evolves complete outcrossing and remains monomorphic otherwise. Darker colours indicate stronger disruptive selection. **B** Evolutionary trajectory in an individual-based simulation. Each point gives the sex allocation (y-axis) of a randomly sampled individual in the population at a particular time (x-axis). The purple shade gives the pollen export rate of that individual, with darker colours indicating higher values. We see that the population first converges to an intermediate sex allocation and pollen export strategy, and then gradually splits into two types: a female-biased hermaphroditic type that does not export any pollen (lighter branch) and a pure male type that exports all its pollen (darker branch). Parameter values used: *c* = 0.75, δ = 0.25, *N* = 10^4^, *µ* = 10^−3^, *σ* = 10^−2^. **C-D-E** Predicted sex allocation (**C**), selfing rate (**D**) and frequency (**E**), of the hermaphroditic type at evolutionary equilibrium, as a function of the cost of pollen dispersal (*c*) and inbreeding depression (δ).

We ran numerical analyses as well as individual-based simulations to investigate the nature of polymorphism evolving after evolutionary branching (Appendix A.1.3.2 for details). These analyses reveal that evolutionary dynamics lead to the coexistence of two highly differentiated types (Fig. 1B): a male type that produces only pollen and exports all of it (i.e., expressing *z*_1_ = 0 and *z*_2_ = 1), and a hermaphroditic type that specialises into ovule production and self-fertilisation, producing little pollen and exporting none of it (*z*_1_ *>* 1*/*2 and *z*_2_ = 0). The hermaphroditic type is heavily female-biased (high *z*_1_, Fig. 1C) and self-fertilises at a rate that varies depending on how much pollen it receives from males, from about 0.25 when the cost of pollen dispersal is low or inbreeding depression is close to *c/*2, to almost 1 when the cost of dispersal is high and inbreeding depression is low (Fig. 1D). This type is always the most frequent in the population, which is due to the asymmetry in transmission created by selfing: males can only ever produce one male offspring per ovule fertilised on a hermaphrodite, whereas hermaphrodites not only produce one hermaphroditic offspring per outcrossed ovule, but also produce two per selfed ovule (Fig. 1E). Thus, the joint evolution of sex allocation and pollen export leads to the emergence of a form of androdioecy where a male type exporting large amounts of pollen persists in the population by fertilising a fraction of the ovules produced by a hermaphroditic type, which is seeking to self-fertilise as many of its ovules as possible. This polymorphism is evolutionarily stable because pollen from different sources compete on equal grounds for fertilisation under classical mass-action, so that hermaphrodites cannot prevent being fertilised by males. We relax this assumption in the next section.

## 4 Maternal control of pollen competition and the rise and fall of sexual specialisation

We now allow maternal plants to manipulate the outcome of pollen competition. Specifically, we let the self-pollen advantage *z*_3_ evolve jointly with sex allocation (*z*_1_) and pollen export (*z*_2_). The resulting selection gradients, derived in Appendix A.3, are given by

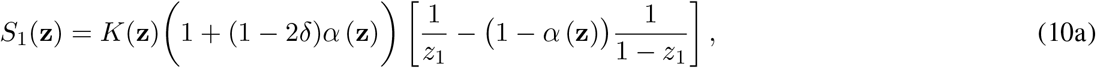

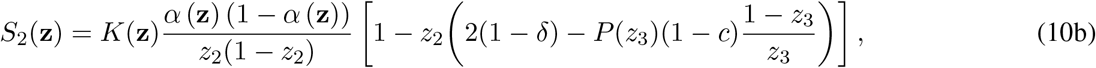

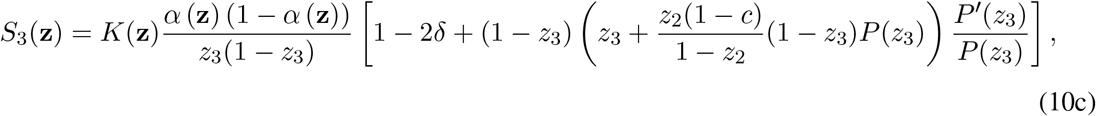

where *P*′(*z*_3_) denotes the first derivative of the pollen export efficiency function *P* (*z*_3_).

Eq. (10a) shows that directional selection on sex allocation *z*_1_ remains unchanged when *z*_3_ evolves jointly with the other traits (compare this equation with eq. (7a) in the previous section), so the singular sex allocation strategy is still given by the one shown in eq. (8b). By contrast, eqs. (10b) and (10c) show that selection gradients on pollen export rate *z*_2_ and the self-pollen advantage *z*_3_ depend on each other in a complicated way. To understand how selection shapes the joint evolution of these two traits, we first consider the simpler case in which self-pollen promotion does not cause pollen discounting (*P* (*z*_3_) = 1 for all *z*_3_).

### 4.1 Cost-free manipulation of pollen competition leads to pure selfing or outcrossing

Assuming *P* (*z*_3_) = 1 for all *z*_3_, the selection gradients on pollen export and self-pollen advantage reduce to

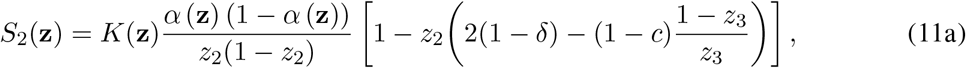

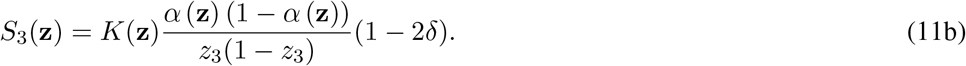

The joint analysis of these two gradients reveals two possible outcomes of directional selection that are determined by the sign of selection on the self-pollen advantage (eq. 11b): either (i) inbreeding depression is strong relative to the transmission advantage (δ *>* 1*/*2) and the population evolves to a complete out-crossing equilibrium where *z*_3_ = 0 (self-pollen cannot fertilise ovules), at which point selection favours complete pollen export (*z*_2_ = 1), and equal allocation male and female function (*z*_1_ = 1*/*2); or (ii) inbreeding depression is weak (δ *<* 1*/*2) and the population evolves towards a complete selfing equilibrium where *z*_3_ = 1 (so that only self-pollen can fertilise ovules), in which case pollen export *z*_2_ becomes effectively neutral as no fitness can be gained as a pollen donor and negligible pollen is needed for selfing (i.e., *S*_2_(**z**) = 0 when *z*_3_ = 1 so that any *z*_2_ *<* 1 is stable, see Appendix A.3.1 for details), and selection favours a female-biased sex allocation as given by eq. (8b). Thus, there can be no mixed mating equilibrium when maternal plants can manipulate pollen competition with no consequence for their pollen export, as predicted by classical theory on prior selfing evolution in the absence of pollen discounting (Lande and Schemske, 1985).

### 4.2 Pollen discounting can generate a mixed mating equilibrium

Eq. (10c) shows that pollen discounting (*P*′(*z*_3_) *<* 0) generates a strictly negative term that selects against self-fertilisation. This countervailing effect could favour the existence of an intermediate equilibrium self-pollen advantage and hence mixed mating. We show in Appendix A.3.2.1 that complete outcrossing (*z*_2_ = 1 and *z*_3_ = 0) and complete selfing (*z*_2_ *<* 1 and *z*_3_ = 1) remain possible endpoints of gradual evolution with pollen discounting. Because eq. (10) does not yield explicit predictions for the outcome of directional selection, we analysed it numerically after specifying the pollen export efficiency function *P* (*z*_3_), which we assume to be given by

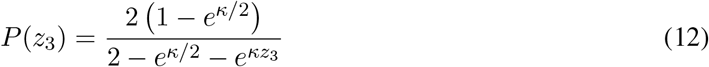

(see Fig. 2A). Equation (12) says *P* (*z*_3_) is a decreasing sigmoid function with *z*_3_ that satisfies *P* (1*/*2) = 1, so that our previous model without maternal control on pollen competition is recovered for *z*_3_ = 1*/*2. The shape of *P* (*z*_3_) is controlled by parameter *κ >* 0. A higher *κ* means stronger sexual interference and so more severe pollen discounting (Fig. 2A).

**Figure 2:**
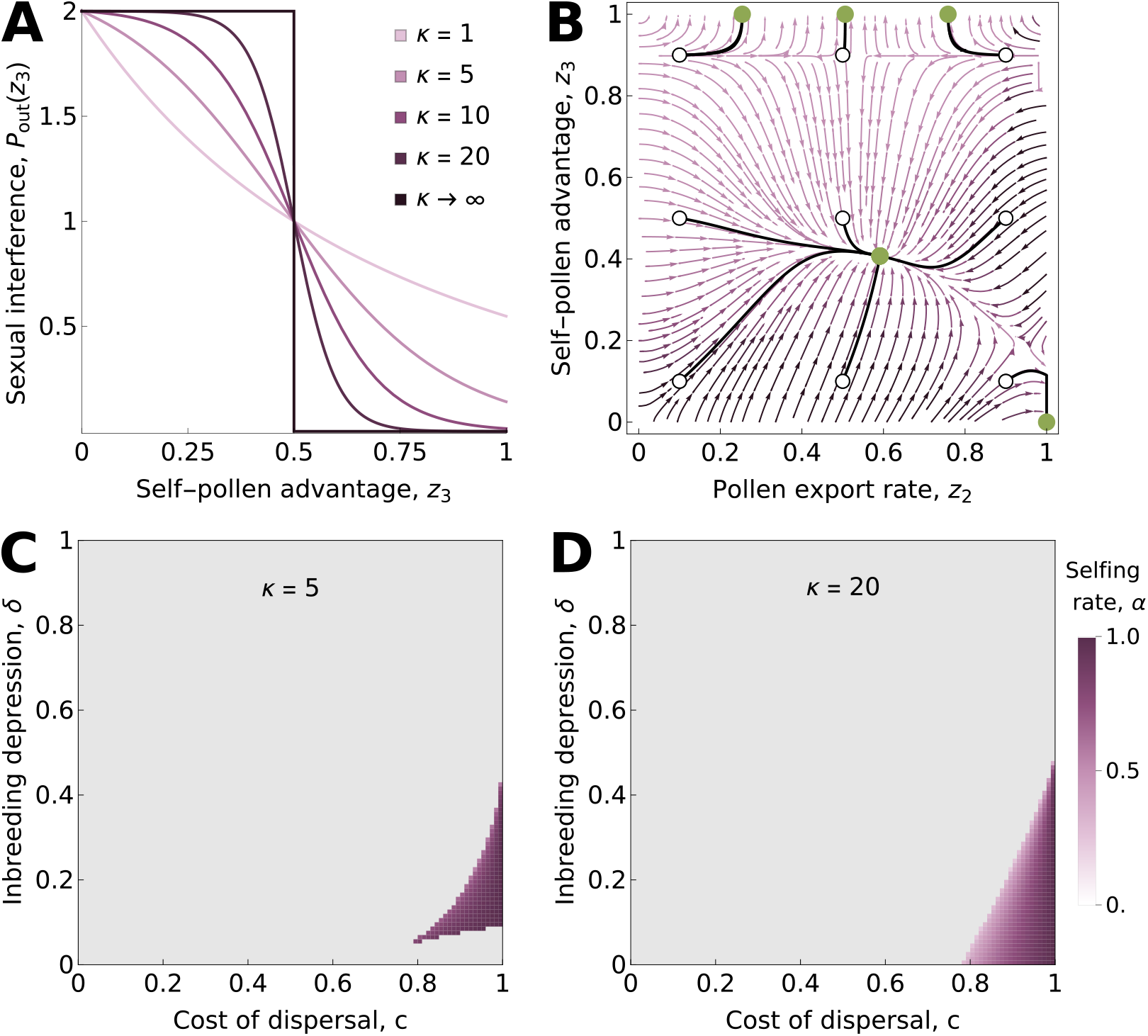
Pollen discounting can generate a mixed-mating equilibrium under maternal control of pollen competition. **A** Pollen export efficiency *P* (*z*_3_) as a function of *z*_3_ for different values of *κ*. Darker shades of purple indicate higher values of *κ* and so stronger sexual interference. **B** Deterministic evolutionary trajectories of pollen export rate *z*_2_ and self-pollen advantage *z*_3_ under directional selection, as inferred from a numerical analysis. The arrows indicate the direction of selection for a given point in the (*z*_2_, *z*_3_) plane, and their colour indicates the intensity of selection with darker shades indicating stronger selection. White points correspond to the nine starting points used in our numerical analyses, solid black lines show the inferred trajectories from these starting points and green points show the attained equilibria. Trajectories reach one of three equilibria: complete outcrossing (bottom right corner), mixed mating (intermediate point) and complete selfing (top edge). The algorithm stops for different values of *z*_2_ in the complete selfing case because the selection gradient on pollen export is zero once self-pollen advantage is *z*_3_ = 1. Parameters used: *κ* = 10, *c* = 0.9, δ = 0.05. **C-D** Selfing rate at the mixed mating equilibrium, where it exists, for moderate (*κ* = 5) and strong sexual interference (*κ* = 20) as a function of the cost of pollen dispersal *c* and inbreeding depression δ. Coloured tiles indicate the selfing rate at the mixed mating equilibrium, and the grey area indicates the parameter range in which no such equilibrium was found.

In addition to the complete selfing and complete outcrossing equilibria, pollen discounting can generate an equilibrium at which the population shows mixed mating (Fig. 2B). The existence of such an equilibrium requires a high cost of pollen dispersal and substantial pollen discounting (high *c* and *κ*). Where it exists, the selfing rate increases with the cost of pollen dispersal *c* and decreases with inbreeding depression δ. Stronger pollen discounting produces a broader range of equilibrium selfing rates (Fig. 2C-D). At this mixed mating equilibrium, the population expresses a female-biased sex allocation given by eq. (8b).

### 4.3 Evolutionary branching leads to complete selfing through transient androdioecy

As shown in section 4.2, directional selection may lead the population to three possible equilibria under pollen discounting: complete selfing, complete outcrossing, or a mixed-mating equilibrium, depending on parameters and initial conditions. Complete selfing and complete outcrossing are endpoints of gradual evolution (see Appendix A.3.2.1 for details), but at the mixed-mating equilibrium, second-order effects of selection may potentially lead to evolutionary branching (since the selection gradient on each trait is zero at this equilibrium).

To study this, we computed **H** (**z**^∗^) numerically at the mixed mating equilibrium for a wide range of parameter combinations. For all of these, we found that **H** (**z**^∗^) has a positive eigenvalue, indicating that selection is disruptive and favours evolutionary branching. Further analysis of **H**(**z**^∗^) reveals that evolutionary branching favours the same kind of polymorphism as before: it favours the emergence of a hermaphroditic type specialising in the production of self-fertilised ovules, and a male type specialising in pollen export. Here, however, the types also diverge in self-pollen advantage: the hermaphroditic type evolves a higher value, which increases its selfing rate, whereas the male evolves a lower value, which improves its pollen export. As the hermaphroditic type evolves an increasingly high self-pollen advantage, fertilisation of its ovules by the male type becomes increasingly unlikely. Male reproductive success and frequency therefore decline until the male type goes extinct (Fig. 3A).

**Figure 3:**
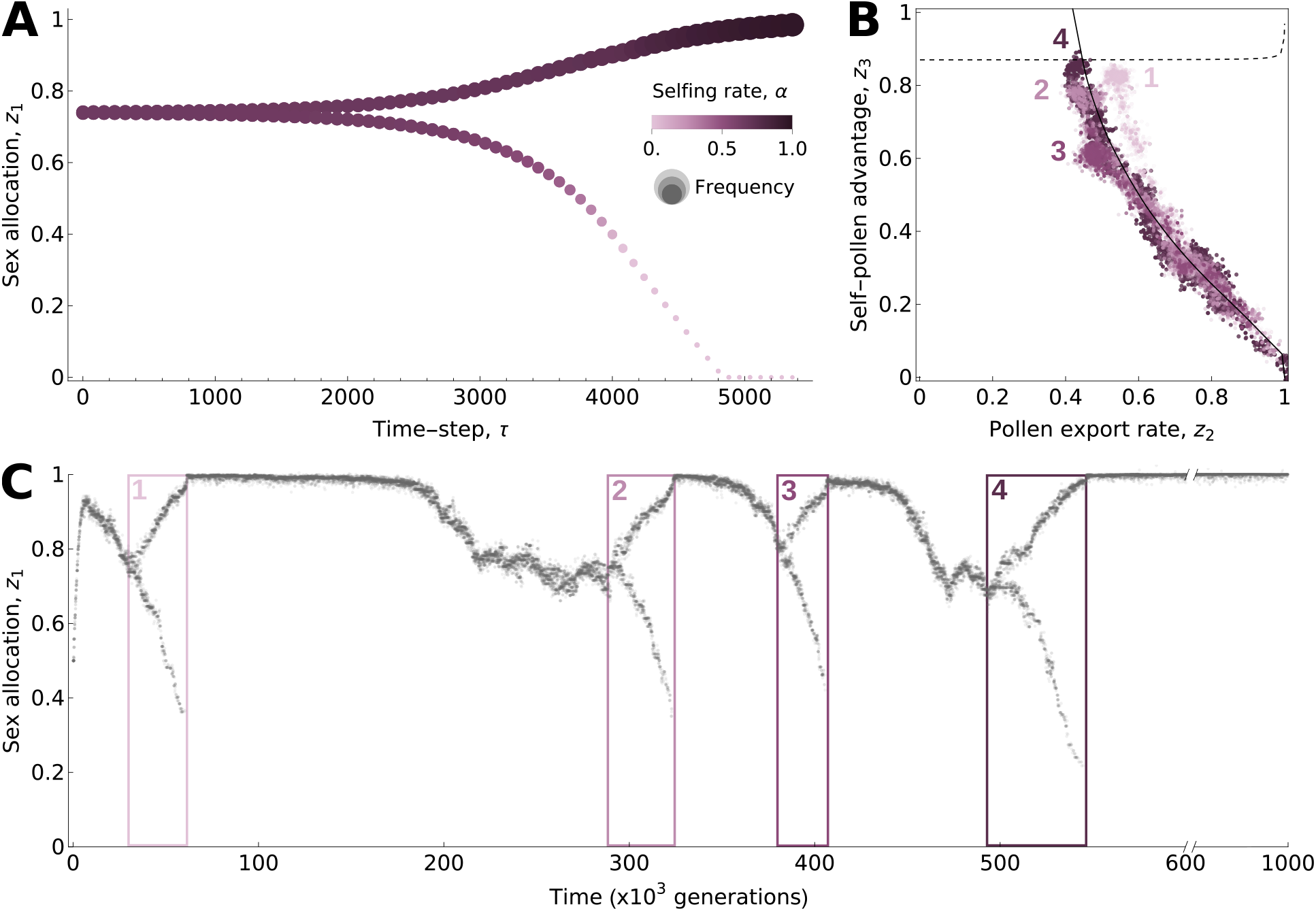
Evolutionary branching produces transient androdioecy and ultimately complete selfing. **A** Deterministic evolutionary trajectory of sex allocation following branching, as inferred by a numerical analysis. Point size indicates the frequency of the type and the shade of purple indicates its selfing rate. As the two types diverge, the selfing rate of the hermaphroditic type increases and the frequency of the male type decreases. **B** Evolutionary trajectory in the (*z*_2_, *z*_3_) phenotypic plane. The solid line gives the predicted trajectory from the numerical analysis (as shown in **A**) and the dotted line gives the threshold self-pollen advantage value above which directional selection favours complete selfing in a monomorphic population. If the male type goes extinct when the hermaphroditic type is below this line, the population evolves back to the mixed mating equilibrium and undergoes evolutionary branching again. Otherwise, the population evolves towards complete selfing. The points show the trajectories followed by the four branching events that took place until complete selfing evolved in the individual-based simulation presented in **C**. The points are coloured per branching event. **C** An example of a simulation where the population undergoes evolutionary branching and evolves back to the mixed equilibrium three times before eventually evolving complete selfing. Parameter set used in all plots: *κ* = 5, *c* = 0.95, δ = 0.25. Other parameters for simulations: *N* = 5 *×* 10^3^, *µ* = 10^−3^, *σ* = 10^−2^.

Once the male type disappears, the population becomes monomorphic for the hermaphroditic type and therefore again evolves under directional selection. Its subsequent trajectory depends on the hermaphroditic phenotype when the male type disappears. If the male type goes extinct when the self-pollen advantage of the hermaphroditic type lies outside the basin of attraction of the complete selfing equilibrium (i.e., sufficiently far from 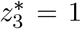; below the dotted line in Fig. 3B), then the population is driven back to the mixed mating equilibrium, at which point disruptive selection again leads to evolutionary branching. If the hermaphroditic type has evolved a high enough self-pollen advantage by the time males disappear, on the other hand, the population evolves towards complete selfing 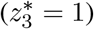 and gradual evolutionary dynamics grind to a halt.

In stochastic individual-based simulations, both outcomes may be observed with probabilities that depend on the parameter values, which determine the strength of disruptive selection and the rate at which the two types diverge, and on population size, which determines how long the male type is expected to persist at low frequencies. However, because complete selfing is an evolutionary endpoint (whereas mixed mating never is in this model), repeated cycles of evolutionary branching and male extinction eventually drive the population to complete selfing (Fig. 3B-C).

## 5 Discussion

Selfing strongly influences sex allocation evolution, yet most theoretical studies treat the mating system as fixed. Models of selfing evolution likewise seldom consider the evolution of sex allocation. In the only study to address their joint evolution, Zhang (2000) modelled the joint evolution of sex allocation and pollen export under a mass-action model of self-fertilisation (Holsinger, 1991). They showed that directional selection on the pollen export rate should be independent of sex allocation, suggesting that selfing influences the evolution of sex allocation but is not influenced in return. Here, we show that selfing and sex allocation do in fact influence the evolution of one another: specifically, correlational selection favours associations between sex allocation and mating-system traits that can generate polymorphic sexual systems.

Under the classical mass-action model, the joint evolution of sex allocation and pollen export leads to a form of androdioecy where hermaphrodites that specialise in the production of self-fertilised ovules co-exist with males that specialise into pollen export. To our knowledge, androdioecy where hermaphroditic individuals do not export any male gamete has not been observed in any plant species. On the other hand, it appears to be more common in animals, notably in nematodes (e.g., in *Caenorhabditis elegans*, Denver et al., 2011) and branchiopod crustaceans (Weeks, 2012), where males coexist with hermaphroditic individuals that do not transfer any sperm (see Chap. 1 in Leonard, 2018 for a review). Our findings thus reveal a possible path from simultaneous hermaphroditism to androdioecy through the joint evolution of sexual and mating strategies that could help explain the evolution of this rare sexual system in some of these animals. However, in most known cases of androdioecy in animals, phylogenetic evidence indicates that this sexual systems evolved from dioecy, probably in response to selection for reproductive assurance in females (Denver et al., 2011; Weeks, 2012; Leonard, 2018). Furthermore, the androdioecy that emerges from the joint evolution of sex allocation and pollen export here is only stable because hermaphrodites cannot avoid fertilisation by males. Androdioecy is never stable once this constraint is lifted, as we showed in our extended model.

When individuals can manipulate gamete competition in favour of their own, we found that the joint evolution of pollen export and the self-pollen advantage makes the existence of a mixed mating equilibrium less likely under directional selection, because prior selfing is now able to evolve where constraints prevented it to do so under classical mass-action. Where a mixed mating equilibrium is reached, evolutionary branching again occurs, leading to the emergence of self-fertilising hermaphrodites and males. However, this polymorphism is no longer stable because hermaphrodites progressively exclude male pollen by evolving a higher self-pollen advantage, causing the extinction of males and thereby driving the evolution of complete selfing through the emergence and collapse of androdioecy. These results show that, contrary to when selfing evolves in isolation (Johnston, 1998; Porcher and Lande, 2005), pollen discounting is not sufficient to ensure the long-term maintenance of mixed mating when sex allocation evolves as well. Moreover, these results expose a limitation of the mass-action model for explaining the widespread occurrence of mixed mating, despite its having sometimes been used for this purpose (e.g., Holsinger, 1991; Porcher and Lande, 2005). Under mass-action, individuals cannot avoid fertilisation by pollen from other individuals, so some outcrossing is unavoidable whenever pollen is exchanged. Mixed mating is therefore built into the model rather than maintained by selection. This highlights the importance of including prior selfing in models seeking to explain the maintenance of mixed mating, especially since this form of selfing is straightforward to achieve mechanistically (Goodwillie et al., 2005).

In our model, mixed mating is unstable because correlated changes in sex allocation and mating-system traits produce increasing fitness returns that favour specialisation and evolutionary branching. Ecological processes that generate diminishing returns to sexual specialisation could oppose this effect. A likely candidate is limited dispersal of pollen and seeds (Fromhage and Kokko, 2010). Limited seed dispersal causes related seeds to compete with one another for recruitment. As investment in female function rises, competition among a plant’s related seeds intensifies, reducing expected fitness returns per seed and hence the benefit of female specialisation (“local resource competition”; Chap. 8 in Maynard Smith, 1978, Clark, 1978; Charnov, 1982; Lloyd and Bawa, 1984; Fromhage and Kokko, 2010; Biernaskie, 2010; Lesaffre et al., 2025). Limited pollen dispersal leads to a similar effect on fitness returns per pollen grain and therefore selects against male specialisation (“local mate/sperm competition”; Hamilton, 1967; Schärer, 2009; Fromhage and Kokko, 2010; Lesaffre et al., 2025). Together, limited dispersal of pollen and seeds could stabilise mixed mating by opposing evolutionary branching in sex allocation. Thus, mechanisms that maintain hermaphroditism may also stabilise mixed mating. Beyond its effect on branching and sex allocation, limited dispersal may favour intermediate selfing rates on its own (Ronfort and Couvet, 1995), but the underlying mechanisms require further investigation.

We assumed that all traits are encoded by a single pleiotropic locus and that individuals are haploid, thereby eliminating potential genetic constraints on phenotypic evolution. Such constraints may oppose evolutionary branching by hindering the build-up of the phenotypic associations favoured by correlational selection, and could thus prevent the emergence of self-fertilising hermaphrodites and males. However, although genetic constraints may pose a strong barrier to the emergence of polymorphism in the short-term, selection may resolve them in the long-term. Selection favouring phenotypic polymorphism can itself reshape genetic architecture and build the trait associations favoured by correlational selection (Rueffler et al., 2006; van Doorn and Dieckmann, 2006; Kopp and Hermisson, 2006; Yeaman and Whitlock, 2011), a process explored in detail by Lesaffre et al. (2025) in the case of the joint evolution of sex allocation and a trait with opposing effects on male and female components of individual fitness. In the biological scenario investigated here, once a mixed-mating equilibrium exists, selection favours evolutionary branching when inbreeding depression is sufficiently weak relative to the cost of pollen dispersal, which is also a condition for the equilibrium to exist. Correlational selection promoting evolutionary branching is therefore expected to persist for as long as the conditions for a mixed mating equilibrium are met, thereby giving the genetic architecture time to evolve.

In the mating model proposed here, the pollen export rate *z*_2_ determines the absolute amount of self-pollen that remains on the plant for selfing, and the self-pollen advantage *z*_3_ determines whether self-pollen grains are at a (dis)advantage relative to cross-pollen grains in the race to ovules. We assumed throughout that these two traits were the target of independent mutations, so that any correlation between them was free to evolve under the action of selection. Biologically, however, changes to floral morphology might sometimes affect both traits simultaneously. For example, reducing herkogamy could increase the absolute amount of self-pollen deposited on a plant’s stigma (decreasing *z*_2_) in addition to causing it to gain a head-start by being deposited earlier (increasing *z*_3_). In this case, mutations modifying the degree of herkogamy would affect both *z*_2_ and *z*_3_ and generate a negative correlation between the two traits due to mutational constraints. Such a negative correlation would accelerate the evolutionary dynamics described here, as it aligns with the direction of correlation selection. Conversely, a positive correlation would slow down the response to selection (Arnold et al., 2008). This reinforces the notion that valuable insights into the joint evolution of sexual and mating systems could be gained from incorporating the details of floral morphology into theoretical models.

Treating inbreeding depression as fixed is conservative with respect to our conclusion that mixed mating is evolutionarily unstable. The mutation load that underlies inbreeding depression is expected to evolve with the selfing rate, but the main effect of selfing on the load is to improve purging, causing inbreeding depression – and so the cost of selfing – to decrease as the selfing rate increases (Lande and Schemske, 1985; Charlesworth et al., 1990, 1991; Abu Awad and Roze, 2020). Purging therefore tends to make the maintenance of mixed mating less likely because it creates a positive feedback loop towards complete selfing, which means that assuming a fixed level of inbreeding depression gives mixed mating a better shot at persisting. More importantly, the correlational selection effect that we describe between sex allocation and mating-system traits does not rely on this assumption: it only requires inbreeding depression to be weak enough for selfing to be net beneficial to maternal parents. We therefore expect that accounting for the dynamics of the load would not fundamentally alter the insights obtained here. Interestingly, recent empirical evidence in hermaphroditic snails suggests that recessive deleterious mutations may not be the only source of inbreeding depression, as epigenetic marks were shown to generate a sizeable reduction in inbred fitness (Bonel et al., 2026). These marks accumulate rapidly and could therefore form the basis of a component of inbreeding depression that is less readily eliminated by purging.

In summary, our results show that the mating and sexual systems of flowering plants may often evolve jointly. We found that this joint evolution can lead to evolutionarily stable androdioecy when plants cannot avoid fertilisation by other individuals. When maternal plants can manipulate the outcome of pollen competition, the joint evolution of selfing and sex allocation ultimately leads to complete selfing through the rise and fall of sexual specialisation. These processes require inbreeding depression to be weak relative to the transmission advantage of selfing. In the future, investigating how additional features of plant ecology, such as pollen limitation or limited dispersal, influence this joint evolution may yield valuable insight into the diversity of plant reproductive strategies.

## Acknowledgements

TL thanks Kai-Hsiu Chen for many inspiring discussions. The authors would also like to thank the Swiss National Science Foundation for funding (SNF grants PZ00-3 232857 to TL, 310030 215135 to JRP and 310030 220139 to CM).

## Statement of Authorship

TL, JRP and CM conceived the study. TL carried out the analytical work and simulations, with input from CM. TL wrote the first draft. All authors contributed to revisions and approved the final manuscript.

## Data and Code Availability

The code used for individual-based simulations, numerical analyses and to check analytical results is available at 10.5281/zenodo.23062245.

## Notes

### Competing Interest Statement

The authors have declared no competing interest.

