## Appendix for "The evolutionary interplay between self-fertilisation and sex allocation in flowering plants"

**B Individual-based simulations** **30**

### Appendix A

### Mathematical and numerical analyses

In this Appendix, we introduce the life cycle followed by individuals in the model and the methodology we use to analyse it. We then provide a detailed account of our analyses.

#### A.1 Methods

##### A.1.1 Life cycle and invasion fitness

Here, we introduce the life cycle that individuals follow. It is presented in a way that does not make any assumption on how traits affect gamete production or mating success through selfing and outcrossing, which allows us to gain general insight into the pathways through which selection acts in this model, irrespective of the particular trait.

We consider a large population composed of $N$ haploid hermaphrodites with discrete generations, where individuals are characterised by their phenotype $\mathbf{z}_i \in \mathbb{R}^V$ for individual $i \in \{1, ..., N\}$, where $V$ is the number of evolving traits. Each generation, individuals pass through the following life-cycle events.

*(i) Sexual development:* First, plants grow and mature sexually, allocating resources to their female and male function according to their phenotype, resulting in the production of $F(\mathbf{z}_i)$ ovules and $M(\mathbf{z}_i)$ pollen grains by plant $i$. Throughout, we assume that the number of ovules is large relative to the number of breeding spots available in the population, so that there always is enough seeds produced to fill these spots and the population remains of constant size $N$. Further, we assume that pollen is much less costly to produce than ovules so that there is always enough pollen for all produced ovules to be fertilised (no pollen limitation).

*(ii) Mating:* Following pollen and ovule production, plants disperse a fraction of their pollen to fertilise

ovules on other plants. Pollen grains received by each plant then compete to fertilise the ovules it carries. The pollen dispersal strategy of individual $i$ and how pollen competition plays out in its pistils depends on its phenotype $\mathbf{z}_i$ as well as the phenotypes of all other individuals in the population, so that individual $i$ self-fertilises at a rate $\alpha(\mathbf{z}_i)$ and the probability that it fertilises an ovule of individual $j \neq i$ is given by $\beta(\mathbf{z}_i, \mathbf{z}_j)$.

*(3) Seed maturation:* Fertilisation results in the production of diploid zygotes. Self-fertilised zygotes suffer from inbreeding depression, becoming viable with probability $1 - \delta$, where $\delta \in [0, 1]$ measures the magnitude of inbreeding depression; meanwhile outcrossed zygotes become viable with probability one. Viable diploid zygotes immediately undergo meiosis to give rise to two haploid seeds each.

*(4) Germination and recruitment:* Following seed maturation, all adults die and the produced seeds germinate and compete for recruitment to fill the $N$ breeding spots in the population.

We assume that individual phenotype $\mathbf{z}_i \in \mathbb{R}^V$ is encoded by a single pleiotropic locus at which alleles may encode any trait combination (or equivalently, $V$ fully linked loci each encoding one trait; "continuum-of-alleles" model, Kimura, 1965; Walsh and Lynch, 2018). Mutations occur at a small fixed rate at this locus, adding small, unbiased and independent deviations onto the trait values encoded by the mutated copy. Under these assumptions, evolution proceeds gradually and its dynamics can be inferred using an adaptive dynamics approach (Avila and Mullon, 2023).

This approach leverages population genetics results to infer the long-term evolutionary trajectory of a population for traits subject to rare mutations with small, unbiased phenotypic effects. The fundamental basis of the adaptive dynamics approach is the *invasion fitness* of a mutant encoding phenotype $\mathbf{z}_\bullet \in \mathbb{R}^V$ arising as a single copy in a resident population fixed for $\mathbf{z} \in \mathbb{R}^V$ at its ecological equilibrium, $W(\mathbf{z}_\bullet, \mathbf{z})$. In a homogeneous population, invasion fitness can be obtained by computing the number of mutant gene copies contributed by a rare focal mutant to the next generation. Under the life cycle presented above, the invasion fitness of a mutant $\mathbf{z}_\bullet$ arising as a single copy in a population otherwise fixed for $\mathbf{z}$ is given by

$$W(\mathbf{z}_\bullet, \mathbf{z}) = \Bigg[\underbrace{F(\mathbf{z}_\bullet)\Big(2\alpha(\mathbf{z}_\bullet)(1-\delta) + 1 - \alpha(\mathbf{z}_\bullet)\Big)}_{\text{Number of mutant seeds produced from own ovules}} + \underbrace{\sum_{i=1}^{N-1} F(\mathbf{z})\beta(\mathbf{z}_\bullet, \mathbf{z})}_{\substack{\text{Number of mutant seeds produced}\\ \text{by fertilising resident ovules}}}\Bigg] \times k(\mathbf{z}_\bullet, \mathbf{z}), \quad \text{(A1a)}$$

where

$$k(\mathbf{z}_\bullet, \mathbf{z}) = \frac{N}{2(N-1)F(\mathbf{z})\left[1 - \delta\,\alpha(\mathbf{z})\right] + 2F(\mathbf{z}_\bullet)\left[1 - \delta\,\alpha(\mathbf{z}_\bullet)\right]} \quad \text{(A1b)}$$

denotes the probability that a given seed is recruited in the population.

Equation (A1) can be understood as follows. The first term between square brackets gives the number of mutant seeds produced by the mutant through its own ovules. The mutant produces $F(\mathbf{z}_\bullet)$ ovules carrying $\mathbf{z}_\bullet$. These ovules then get self-fertilised in proportion $\alpha(\mathbf{z}_\bullet)$, resulting in selfed zygotes that become viable with probability $1-\delta$ and divide into two mutant haploid seeds if they do so. The rest of the ovules $(1-\alpha(\mathbf{z}_\bullet))$ gets fertilised by resident pollen, resulting in outcrossed, heterozygous zygotes that become viable with probability 1 and subsequently divide to produce one mutant seed and one resident seed. This reduced diploid phase therefore captures the transmission advantage of self-fertilisation, which is diminished by inbreeding depression (Fisher, 1941; Lande and Schemske, 1985). The second term between square brackets gives the number of mutant seeds that the mutant produces by fertilising ovules on resident individuals. Each of the $N-1$ resident individuals in the population produces $F(\mathbf{z})$ ovules, a fraction $\beta(\mathbf{z}_\bullet, \mathbf{z})$ of which gets fertilised by the mutant. Thus, the term between square brackets on the first line of eq. (A1) gives the number of mutant seeds produced by the focal mutant through all reproductive pathways. The quantity $k(\mathbf{z}_\bullet, \mathbf{z})$ gives the probability that a seed gets recruited in the population in the next time-step, which is obtained as the number of available breeding spots ($N$) divided by the total number of seeds produced by the population.

In the limit where the population becomes large ($N \to \infty$), eq. (A1a) simplifies to

$$W(\mathbf{z}_\bullet, \mathbf{z}) = K(\mathbf{z})\left[\Big(1 + \alpha(\mathbf{z}_\bullet)(1-2\delta)\Big)\frac{F(\mathbf{z}_\bullet)}{F(\mathbf{z})} + B(\mathbf{z}_\bullet, \mathbf{z})\right], \quad \text{(A1c)}$$

where

$$K(\mathbf{z}) = \lim_{N\to\infty} F(\mathbf{z})k(\mathbf{z}_\bullet, \mathbf{z}) = \frac{1}{2\left[1-\delta\,\alpha\left(\mathbf{z}\right)\right]} \tag{A1d}$$

is a strictly positive term that captures the effect of density regulation, and

$$B(\mathbf{z}_\bullet, \mathbf{z}) = \lim_{N\to\infty} \left( \sum_{i=1}^{N-1} \beta\left(\mathbf{z}_\bullet, \mathbf{z}\right) \right) \tag{A1e}$$

denotes the mutant's reproductive success as a pollen donor. Equation (A1c) corresponds to eq. (3) in the main text, and is used for all subsequent analyses.

##### A.1.2 Gradual evolution in a monomorphic population

Under the assumption that mutants are rare and have small, unbiased phenotypic effects, the gradual evolutionary dynamics of the population can be inferred by studying invasion fitness $W(\mathbf{z}_\bullet, \mathbf{z})$, and can be decomposed into two phases.

###### A.1.2.1 Directional selection

At first, a population monomorphic for some phenotype $\mathbf{z}$ will experience directional selection. The direction and strength of selection on a given trait $z_v$, $v \in \{1, ..., V\}$, during this phase is given by the linear selection gradient on this trait,

$$S_v(\mathbf{z}) = \left.\frac{\partial W(\mathbf{z}_\bullet, \mathbf{z})}{\partial z_{v\bullet}}\right|_\circ, \tag{A2a}$$

where the $|_\circ$ notation indicates that the derivative is evaluated at $\mathbf{z}_\bullet = \mathbf{z}$. A positive gradient means that alleles encoding a higher value of $z_v$ are favoured by natural selection, whereas a negative gradient indicates that alleles encoding a lower value are favoured. In our model, this is

$$S_v(\mathbf{z}) = K(\mathbf{z}) \left[ \frac{1}{F(\mathbf{z})} \left.\frac{\partial F(\mathbf{z}_\bullet)}{\partial z_{v\bullet}}\right|_\circ \left(1+\alpha\left(\mathbf{z}\right)\left(1-2\delta\right)\right) + \left(1-2\delta\right) \left.\frac{\partial \alpha\left(\mathbf{z}_\bullet\right)}{\partial z_{v\bullet}}\right|_\circ + \left.\frac{\partial B(\mathbf{z}_\bullet, \mathbf{z})}{\partial z_{v\bullet}}\right|_\circ \right]. \tag{A2b}$$

Equation (A2b) reveals that directional selection on a trait $z_v \in \mathbf{z}$ acts through three distinct pathways. The first pathway corresponds to the fitness effect of a change in $z_v$ through its effect on ovule production. The term $\frac{1}{F(\mathbf{z})} \left.\frac{\partial F(\mathbf{z}_\bullet)}{\partial z_{v\bullet}}\right|_\circ$ gives the effect of a small change in $z_v$ on the relative production of ovules, meanwhile the term between brackets $1 + \alpha(\mathbf{z})(1 - 2\delta)$ gives the average number of mutant copies transmitted per ovule by the mutant for a given selfing rate $\alpha(\mathbf{z})$. The product of these two terms therefore gives the relative change in the number of mutant copies transmitted by the mutant through its own ovules, for a selfing rate $\alpha(\mathbf{z})$. The second pathway is the fitness effect of a change in $z_v$ through its effect on the selfing rate. When inbreeding depression is low ($\delta < 1/2$), self-fertilised seeds transmit on average more mutant copies than the outcrossed and so selection will favour trait values that increase the selfing rate through this component. For example, if an increase in trait $z_v$ results in a decrease in selfing rate ($\left.\frac{\partial \alpha(\mathbf{z}_\bullet)}{\partial z_{v\bullet}}\right|_\circ < 0$) then this effect favours lower values of $z_v$ (the second component is negative). Conversely, when inbreeding depression is high ($\delta > 1/2$), outcrossed seeds transmit more mutant copies on average, so that selection favours trait values that decrease the selfing rate. The third and last pathway is the fitness effect of a change in $z_v$ through its effect on its success as a pollen donor.

Through the sequential fixation of mutants of small effect, the population will travel in phenotypic space from one monomorphic state to the next in the direction favoured by selection, which is given by the vector of selection gradients $\mathbf{S}(\mathbf{z}) = \big(\, S_v(\mathbf{z}) \,\big)_{1 \leqslant v \leqslant V}$. Eventually, it may reach a singular phenotype $\mathbf{z}^*$ for which directional selection vanishes†, that is,

$$\mathbf{z}^* \quad \text{such that} \quad \mathbf{S}(\mathbf{z}) = \mathbf{0}. \tag{A3}$$

Formally, whether the population will evolve towards a singular phenotype $\mathbf{z}^*$ can be assessed by computing the Jacobian matrix at this point, $\mathbf{J}(\mathbf{z}^*)$, which is a $V \times V$ matrix the $(u, v)$-entry $J_{uv}(\mathbf{z}^*)$ of which is given by

$$J_{uv}(\mathbf{z}^*) = \left.\frac{\partial S_u(\mathbf{z})}{\partial z_v}\right|_*, \tag{A4}$$

where the notation $|_*$ indicates that the derivative is evaluated at the singular point (i.e., $\mathbf{z}_\bullet = \mathbf{z} = \mathbf{z}^*$).

†We note that in some cases, directional selection may also never vanish and drive a trait against a boundary of its definition domain. For example, for a trait $z_v \in [0, 1]$, selection may always favour a decrease in $z_v$ such that $S_v(\mathbf{z}) < 0$ even once $z_v = 0$. In this case, $z_v = 0$ is stable so long as the population remains monomorphic for all other traits. If other traits undergo evolutionary branching irrespective of $z_v$ (see below), the subsequent evolution of this trait requires additional analysis.

Convergence to the singular point $\mathbf{z}^*$ requires that the leading eigenvalue of $\mathbf{J}\left(\mathbf{z}^*\right)$ is negative under the assumption that all traits mutate independently (Leimar, 2009). In some cases, however, inspecting the selection gradients directly is sufficient to assess convergence stability.

###### A.1.2.2 Numerical analysis of evolution under directional selection

When several traits evolve jointly, explicit mathematical investigations of evolutionary dynamics under directional selection can become difficult. Nevertheless, useful insight can be gained from a numerical analysis where one follows the deterministic trajectory of a monomorphic population in phenotypic space, as dictated by selection gradients. Specifically, given a population monomorphic for phenotype $\mathbf{z}(\tau)$ at some time-step $\tau$, we can compute the vector of selection gradients acting on the traits at that point in phenotypic space, $\mathbf{S}\big(\mathbf{z}(\tau)\big)$, and use it to obtain the phenotype expressed by the population in the next time-step as

$$z_v(\tau+1) = \mathcal{C}_{[z_v^{\min}, z_v^{\max}]}\Big(z_v(\tau) + \Delta\; S_v\big(\mathbf{z}(\tau)\big)\Big) \tag{A5}$$

where $v \in \{1, ...V\}$, $0 < \Delta \ll 1$ is a small deviation and $\mathcal{C}_{[z_v^{\min}, z_v^{\max}]} : \mathbb{R} \to [z_v^{\min}, z_v^{\max}]$ denotes the clipping function that maintains trait $z_v$ within its definition domain:

$$\mathcal{C}_{[z_v^{\min}, z_v^{\max}]}(z_v) = \begin{cases} z_v^{\min} & \text{if} \quad z_v \leqslant z_v^{\min} \\ z_v & \text{if} \quad z_v^{\min} < z_v < z_v^{\max} \\ z_v^{\max} & \text{if} \quad z_v^{\max} \leqslant z_v. \end{cases} \tag{A6}$$

Throughout, we assume that the small deviation $\Delta$ is a random variable

$$\Delta \sim \mathcal{U}(\Delta_{\min}, \Delta_{\max}), \tag{A7}$$

where $\mathcal{U}(x, y)$ denotes the uniform distribution on $[x, y] \subset \mathbb{R}$ and $0 < \Delta_{\min} < \Delta_{\max}$ give the minimal and maximal possible deviations. This ensures that the algorithm does not remain trapped in stable oscillations around its equilibrium.

To determine the equilibrium of directional selection reached by a population initially monomorphic for some phenotype $\mathbf{z}(0)$, we can iterate eq. (A5) for all traits numerically until we reach a phenotype

$$\mathbf{z}^* \quad \text{such that} \quad |z_v(\tau+1) - z_v(\tau)| < \epsilon \quad \text{for all} \quad z_v \in \mathbf{z}, \tag{A8}$$

where $\epsilon > 0$ is a small tolerance criterion.

This algorithm is implemented in the *Mathematica* file named `Numerical_DirectionalSelection.nb`, available at `10.5281/zenodo.23062245`.

###### A.1.2.3 Disruptive selection

Once the population expresses $\mathbf{z}^*$, leading order forces of selection vanish, so that second-order effects come into play. The nature of selection on the traits is then given by the Hessian matrix $\mathbf{H}(\mathbf{z}^*)$, which is a $V \times V$ matrix the $(u, v)$-entry $H_{uv}(\mathbf{z}^*)$ of which is given by

$$H_{uv}(\mathbf{z}^*) = \left. \frac{\partial^2 W(\mathbf{z}_\bullet, \mathbf{z})}{\partial z_{u\bullet}\, \partial z_{v\bullet}} \right|_{\mathbf{z}_\bullet = \mathbf{z} = \mathbf{z}^*}. \tag{A9a}$$

Elements on the diagonal of $\mathbf{H}(\mathbf{z}^*)$ are given by the second derivative of $W(\mathbf{z}_\bullet, \mathbf{z})$ with respect to mutant trait value $H_{vv}(\mathbf{z}^*)$, evaluated at $\mathbf{z}_\bullet = \mathbf{z} = \mathbf{z}^*$, and therefore give the quadratic selection coefficient on $z_v$ at the singular point. This coefficient captures the nature of selection acting on each trait independently. A negative value of $H_{vv}(\mathbf{z}^*)$ indicates that selection on trait $z_v$ is stabilising and so favours a decrease in variance around $z_v^*$. A positive value, meanwhile, indicates that selection on $z_v$ is disruptive at the singular point, and so favours an increase in variance from $z_v^*$. Off-diagonal elements (so where $u \neq v$) are the cross-derivative of $W(\mathbf{z}_\bullet, \mathbf{z})$ with respect to different trait values of the mutant. These elements correspond to the correlational selection coefficient between these two traits. A positive value of $H_{uv}(\mathbf{z}^*)$ indicates that selection favours a positive correlation between the two traits, whereas a negative $H_{uv}(\mathbf{z}^*)$ shows that selection favours a negative correlation. The evolution of a population expressing $\mathbf{z}^*$ is determined by the leading eigenvalue of $\mathbf{H}(\mathbf{z}^*)$. If it is negative, then selection is stabilising in all dimensions of phenotype, favouring the population to remain monomorphic for $\mathbf{z}^*$, so that phenotypic evolution comes to a halt. If the leading eigenvalue is positive, on the other hand, disruptive selection will lead to the gradual emergence of polymorphism via evolutionary branching. It is useful to note here that there is two main ways for the leading eigenvalue of $\mathbf{H}(\mathbf{z}^*)$ to be positive. Either one diagonal element $H_{vv}(\mathbf{z}^*)$ is positive, meaning that one trait will undergo branching irrespective of selection on other traits, which may in turn cause other traits to become polymorphic; or all diagonal elements $H_{vv}(\mathbf{z}^*)$ are negative, so that selection would be stabilising for $\mathbf{z}^*$ if all traits evolved in isolation, but correlational selection amongst traits is sufficiently strong to overcome stabilising selection on each trait and lead to branching (Débarre et al., 2014). Thus, the more traits co-evolve, the more routes to polymorphism there is.

In the context of our model, the $(u, v)$-element of $\mathbf{H}(\mathbf{z}^*)$ associated with traits $z_u$ and $z_v$, $H_{uv}(\mathbf{z}^*)$, is given by

$$\begin{aligned} H_{uv}(\mathbf{z}^*) = K(\mathbf{z}) \times \Bigg[ & \frac{1}{F(\mathbf{z})} \frac{\partial^2 F(\mathbf{z}_\bullet)}{\partial z_{u\bullet} \partial z_{v\bullet}}\bigg|_* \Big(1 + \alpha(\mathbf{z})(1-2\delta)\Big) \\ & + (1-2\delta)\left( \frac{\partial^2 \alpha(\mathbf{z}_\bullet)}{\partial z_{u\bullet}\partial z_{v\bullet}}\bigg|_* + \frac{\partial \alpha(\mathbf{z}_\bullet)}{\partial z_{u\bullet}}\bigg|_* \times \frac{1}{F(\mathbf{z})} \frac{\partial F(\mathbf{z}_\bullet)}{\partial z_{v\bullet}}\bigg|_* + \frac{\partial \alpha(\mathbf{z}_\bullet)}{\partial z_{v\bullet}}\bigg|_* \times \frac{1}{F(\mathbf{z})} \frac{\partial F(\mathbf{z}_\bullet)}{\partial z_{u\bullet}}\bigg|_* \right) \\ & + \frac{\partial^2 B(\mathbf{z}_\bullet, \mathbf{z})}{\partial z_{u\bullet}\partial z_{v\bullet}}\bigg|_* \Bigg]. \end{aligned} \tag{A9b}$$

Equation (A9b) shows that second-order forces of selection act through similar pathways to directional selection. The first term between square brackets captures the fitness effect of a joint change in $z_u$ and $z_v$ via its effect on ovule production, for a given selfing rate. The second term captures the fitness effect of a joint change in $z_u$ and $z_v$ through the change it causes in the number of self-fertilised ovules, which can either result from the joint effect of $z_u$ and $z_v$ on the selfing rate, or from the effect of one trait on ovule production and of the other on the selfing rate. Similar to the directional selection case, the direction of this effect depends on whether outcrossed or selfed seeds yield higher fitness returns, i.e., on whether inbreeding depression is greater or lower than a half, respectively. The last term captures the effect of a joint change in $z_u$ and $z_v$ on reproductive success as a pollen donor.

##### A.1.3 The nature of the polymorphism resulting from evolutionary branching

###### A.1.3.1 Analytical setup

When polymorphism emerges through evolutionary branching, the nature of the polymorphism maintained at equilibrium can be studied by modelling a dimorphic resident population in which two types of individuals, say A and B expressing strategies $\mathbf{z}_\mathrm{A}$ and $\mathbf{z}_\mathrm{B}$, respectively, coexist at an ecological equilibrium. Under the assumption that the population is large ($N \to \infty$), this equilibrium can be described by the equilibrium frequency of type A,

$$\hat{p}(\mathbf{z}_\mathrm{A}, \mathbf{z}_\mathrm{B}) \quad \text{such that} \quad \Delta p(\mathbf{z}_\mathrm{A}, \mathbf{z}_\mathrm{B}) = p_{t+1} - p_t = 0, \tag{A10a}$$

where $p_t$ is the frequency of type A at time $t$ and $\Delta p(\mathbf{z}_\mathrm{A}, \mathbf{z}_\mathrm{B})$ is the change in this frequency over one generation. In our model, the frequency change is given by

$$\begin{aligned}\Delta p(\mathbf{z}_\mathrm{A}, \mathbf{z}_\mathrm{B}) = & \frac{2F(\mathbf{z}_\mathrm{A})p_t}{\overline{F}(\mathbf{z}_\mathrm{A}, \mathbf{z}_\mathrm{B})}\left[\alpha\left(\mathbf{z}_\mathrm{A}\right)(1-\delta) + \beta\left(\mathbf{z}_\mathrm{A}, \mathbf{z}_\mathrm{A}\right)p_t + \frac{1}{2}\beta\left(\mathbf{z}_\mathrm{B}, \mathbf{z}_\mathrm{A}\right)(1-p_t)\right] \\ & + \frac{2F(\mathbf{z}_\mathrm{B})(1-p_t)}{\overline{F}(\mathbf{z}_\mathrm{A}, \mathbf{z}_\mathrm{B})}\frac{1}{2}\beta\left(\mathbf{z}_\mathrm{A}, \mathbf{z}_\mathrm{B}\right)p_t \; - \; p_t,\end{aligned} \tag{A10b}$$

where

$$\overline{F}(\mathbf{z}_\mathrm{A}, \mathbf{z}_\mathrm{B}) = 2F(\mathbf{z}_\mathrm{A})\left[1 - \delta\,\alpha\left(\mathbf{z}_\mathrm{A}\right)\right]p_t + 2F(\mathbf{z}_\mathrm{B})\left[1 - \delta\,\alpha\left(\mathbf{z}_\mathrm{B}\right)\right](1-p_t) \tag{A10c}$$

is the average number of mature seeds produced per individual in the population at time $t$.

The term on the first line in eq. (A10b) represents the proportion of the next generation that carries $\mathbf{z}_\mathrm{A}$ and has a type A individual as maternal parent. The ratio on the left-hand side gives proportion of seeds in the next generation produced by these parents, and the term between square brackets gives the probability that these seeds carry $\mathbf{z}_\mathrm{A}$ depending on how they were fertilised. Each type A individual self-fertilises at rate $\alpha\left(\mathbf{z}_\mathrm{A}\right)$, in which case the resulting diploid zygote survives with probability $1-\delta$ and divides to produce two seeds carrying $\mathbf{z}_\mathrm{A}$; a fraction $\beta\left(\mathbf{z}_\mathrm{A}, \mathbf{z}_\mathrm{A}\right)p_t$ of its ovules is fertilised by outcross pollen carrying $\mathbf{z}_\mathrm{A}$, in which case the diploid zygote survives with probability one and yields only seeds carrying $\mathbf{z}_\mathrm{A}$; finally, a fraction $\beta\left(\mathbf{z}_{\mathrm{B}},\mathbf{z}_{\mathrm{A}}\right)(1-p_t)$ of its ovules are fertilised by $\mathbf{z}_{\mathrm{B}}$ pollen and the resulting zygote survives with probability one, dividing to yield one seed carrying $\mathbf{z}_{\mathrm{A}}$ and the other carrying $\mathbf{z}_{\mathrm{B}}$. The term on the second line gives the proportion of the population in the next generation that carries $\mathbf{z}_{\mathrm{A}}$ and has a type B individual as maternal parent. Similar to type A maternal parents, the ratio on the left-hand side gives the proportion of seeds in the next generation produced by these parents. It is multiplied by the probability that these seeds carry $\mathbf{z}_{\mathrm{A}}$, which is only the case for $1/2$ of the seeds produced through outcrossing with $\mathbf{z}_{\mathrm{A}}$-bearing pollen, which occurs in proportion $\beta\left(\mathbf{z}_{\mathrm{A}},\mathbf{z}_{\mathrm{B}}\right)p_t$.

In a polymorphic population at its ecological equilibrium (as given by eq. A10), we now consider the fate of a rare mutant $\mathbf{z}_\bullet$ arising as a single copy, which is again determined by its invasion fitness, denoted as $W_{\mathrm{p}}(\mathbf{z}_\bullet,\mathbf{z}_{\mathrm{A}},\mathbf{z}_{\mathrm{B}})$ to signify that the mutant $\mathbf{z}_\bullet$ invades into a population that is polymorphic for $\mathbf{z}_{\mathrm{A}}$ and $\mathbf{z}_{\mathrm{B}}$. Under the assumption that the mutant deviates weakly of either type A or type B residents, the mutant's invasion fitness can be used to obtain the selection gradient on each resident's phenotype. Specifically, the selection gradient on the value of trait $z_v$ encoded by resident type $\mathrm{x}\in\{\mathrm{A},\mathrm{B}\}$, $z_{v,\mathrm{x}}$ can be obtained as

$$S_{v,\mathrm{x}}(\mathbf{z}_{\mathrm{A}},\mathbf{z}_{\mathrm{B}})=\left.\frac{\partial W_{\mathrm{p}}(\mathbf{z}_\bullet,\mathbf{z}_{\mathrm{A}},\mathbf{z}_{\mathrm{B}})}{\partial z_{v\bullet}}\right|_{\mathbf{z}_\bullet=\mathbf{z}_{\mathrm{x}}}. \tag{A11}$$

Similar to the monomorphic case, a positive selection gradient indicates that selection favours an increase in trait $z_{v,\mathrm{x}}$ from its current value, whereas a negative gradient indicates that a decrease in trait $z_{v,\mathrm{x}}$ is favoured by selection. In this setting, small-effect mutants arising either from the type A or the type B allele will be able to invade and replace their ancestral allele if they encode a phenotype taking their type further in the direction favoured by selection, given by the type-specific vectors of selection gradients $\mathbf{S}_{\mathrm{A}}(\mathbf{z}_{\mathrm{A}},\mathbf{z}_{\mathrm{B}})=\left(\,S_{v,\mathrm{A}}(\mathbf{z}_{\mathrm{A}},\mathbf{z}_{\mathrm{B}})\,\right)_{1\leqslant v\leqslant V}$ and $\mathbf{S}_{\mathrm{B}}(\mathbf{z}_{\mathrm{A}},\mathbf{z}_{\mathrm{B}})=\left(\,S_{v,\mathrm{B}}(\mathbf{z}_{\mathrm{A}},\mathbf{z}_{\mathrm{B}})\,\right)_{1\leqslant v\leqslant V}$.

Eventually, this invasion-and-replacement process may lead the population to an equilibrium coalition where the gradients vanish, i.e.,

$$\mathbf{z}^*_{\mathrm{A}},\mathbf{z}^*_{\mathrm{B}}\quad\text{such that}\quad\mathbf{S}_{\mathrm{A}}(\mathbf{z}^*_{\mathrm{A}},\mathbf{z}^*_{\mathrm{B}})=\mathbf{S}_{\mathrm{B}}(\mathbf{z}^*_{\mathrm{A}},\mathbf{z}^*_{\mathrm{B}})=0, \tag{A12}$$

or push traits against a boundary of their definition domain (e.g., $S_{v,\mathrm{x}}(\mathbf{z}_{\mathrm{A}},\mathbf{z}_{\mathrm{B}})<0$ when $z_{v,\mathrm{x}}=0$), so that evolution owing to directional selection on each type comes to a halt. Determining whether this equilibrium coalition is evolutionarily stable (in the sense that neither of the two branches undergo another round of branching once they sit at an equilibrium of directional selection $\mathbf{z}_{\mathrm{x}}^{*}$) would require computing type-specific Hessian matrices, but we leave this aside here.

###### A.1.3.2 Numerical analyses

It is usually difficult to solve explicitly for the equilibrium polymorphic coalition favoured by natural selection, especially if several traits evolve jointly. We therefore resort to a numerical analysis in which we follow the deterministic trajectories of the phenotypes expressed by type A and B from an arbitrary initial coalition $(\mathbf{z}_{\mathrm{A}}, \mathbf{z}_{\mathrm{B}})$ across phenotypic space, as dictated by the vectors of gradients $\mathbf{S}_{\mathrm{A}}(\mathbf{z}_{\mathrm{A}}, \mathbf{z}_{\mathrm{B}})$ and $\mathbf{S}_{\mathrm{B}}(\mathbf{z}_{\mathrm{A}}, \mathbf{z}_{\mathrm{B}})$. Specifically, given the phenotypes expressed by type A and B in the population at some time-step $\tau$, $\mathbf{z}_{\mathrm{A}}(\tau)$ and $\mathbf{z}_{\mathrm{B}}(\tau)$, the elements $z_{v,\mathrm{x}}(\tau+1)$ of the phenotype $\mathbf{z}_{\mathrm{x}}(\tau+1)$ expressed by type x in the next time-step are obtained as

$$z_{v,\mathrm{x}}(\tau+1) = \mathcal{C}_{[z_v^{\min}, z_v^{\max}]}\Big(z_{v,\mathrm{x}}(\tau) + \Delta\, S_{v,\mathrm{x}}\big(\mathbf{z}_{\mathrm{A}}(\tau), \mathbf{z}_{\mathrm{B}}(\tau)\big)\Big) \tag{A13}$$

where $0 < \Delta \ll 1$ is a small deviation and $\mathcal{C}_{[z_v^{\min}, z_v^{\max}]} : \mathbb{R} \to [z_v^{\min}, z_v^{\max}]$ denotes the clipping function, similar to the monomorphic case described in section A.1.2.2. To avoid oscillatory behaviours, we also assume here that the small deviation $\Delta$ is a random variable

$$\Delta \sim \mathcal{U}(\Delta_{\min}, \Delta_{\max}). \tag{A14}$$

To determine the evolutionary equilibrium reached by a dimorphic population, we can iterate eq. (A13) for all traits numerically until we reach

$$\mathbf{z}_{\mathrm{A}}^{*},\ \mathbf{z}_{\mathrm{B}}^{*} \quad \text{such that} \quad |z_{v,\mathrm{A}}(\tau+1) - z_{v,\mathrm{A}}(\tau)| < \epsilon \quad \text{and} \quad |z_{v,\mathrm{B}}(\tau+1) - z_{v,\mathrm{B}}(\tau)| < \epsilon \tag{A15}$$

for all $z_{v,\mathrm{A}} \in \mathbf{z}_{\mathrm{A}}$ and $z_{v,\mathrm{B}} \in \mathbf{z}_{\mathrm{B}}$, where $\epsilon > 0$ is a small tolerance criterion, similar to the monomorphic case (Section A.1.2.2); or until one type displaces the other (i.e., until the equilibrium frequency in the population defined by eq. (A10a) hits $\hat{p}(\mathbf{z}_\mathrm{A}, \mathbf{z}_\mathrm{B}) = 0$ or 1), at which point polymorphism is lost so the population is again monomorphic and evolving under directional selection.

There is a couple of options for how to initialise the numerical recursion described above. A first option is to compute the leading eigenvector of the Hessian matrix $\mathbf{H}(\mathbf{z}^*)$ (eq. A9a) at the singular point, $\boldsymbol{\eta}(\mathbf{z}^*)$. This vector gives the direction in which branching initially takes place in phenotypic space. We may then position the two types along this vector in the close vicinity of the singular point $\mathbf{z}^*$, i.e.,

$$\mathbf{z}_\mathrm{A}(\tau = 0) = \mathbf{z}^* + \epsilon\, \boldsymbol{\eta}(\mathbf{z}^*) \quad \text{and} \quad \mathbf{z}_\mathrm{B}(\tau = 0) = \mathbf{z}^* - \epsilon\, \boldsymbol{\eta}(\mathbf{z}^*), \tag{A16}$$

where $0 < \epsilon \ll 1$ is a small constant. Iterating eq. (A13) for all traits from this starting point gives the deterministic trajectory that evolutionary branching is expected to follow from this singular point.

This first option works for most of the parameter space. However, for some parameter values for which selection is fairly weak, the two types may take a very long time to diverge and reach their respective equilibrium. To circumvent this issue and save computation time, a second option is to start the algorithm with strongly diverged types in the direction given by the eigenvector $\boldsymbol{\eta}(\mathbf{z}^*)$, provided that such polymorphic situation is possible (both types are maintained in a stable coalition). The algorithm then rapidly converges to an equilibrium, because greater phenotypic differences between types tend amplify selection when the branches have not yet reached their respective equilibria. We ran the algorithm using both initialisations whenever possible for our models, and found that the same equilibrium was always reached.

| |
|---|
| This algorithm is implemented in the *Mathematica* file named `Numerical_DisruptiveSelection.nb`, available at `10.5281/zenodo.23062245`. |

#### A.2 The joint evolution of sex allocation and pollen export

In this section, we derive the analytical results presented in Section 3 of the main text. We consider the joint evolution of two traits, sex allocation $z_1 \in [0,1]$, which determines the proportion of its reproductive resources that an individual commits to its female function, and pollen export $z_2 \in [0,1]$, which determines the proportion of its pollen that an individual disperses to fertilise ovules on other plants. The population follows the life cycle described in Section A.1.1, and the two traits affect the behaviour of individuals along this life cycle as follows.

*(i) Sexual development:* The number of ovules and pollen grains produced by individual $i$, $F(\mathbf{z}_i)$ and $M(\mathbf{z}_i)$ are given by

$$F(\mathbf{z}_i) = F_0 z_{1,i} \quad \text{and} \quad M(\mathbf{z}_i) = M_0(1 - z_{1,i}), \tag{A17}$$

respectively, where $F_0 \gg 0$ is the maximum female fecundity (the number of ovules produced by an individual allocating all its resources to its female function) and $M_0 \gg F_0$ is the maximum male fecundity. Thus, ovule and pollen production depend linearly on resource allocation, and pollen export does not influence gamete production.

*(ii) Mating:* After sexual development, plant $i$ disperses a fraction $z_{2,i}$ of its pollen to fertilise ovules on other plants, and retains the remainder fraction $1 - z_{2,i}$ for self-fertilisation, following a mass-action model (Holsinger, 1991; Porcher and Lande, 2005). A fraction $c \in [0,1]$ of the dispersed pollen is lost during dispersal, and the successful fraction $1 - c$ is distributed equally amongst the $N - 1$ other plants. As a result, the selfing rate $\alpha(\mathbf{z}_i)$ of individual $i$ is given by the proportion of its own pollen out of the total amount of pollen it received, i.e.,

$$\alpha(\mathbf{z}_i) = \frac{M(\mathbf{z}_i)(1 - z_{2,i})}{M(\mathbf{z}_i)(1 - z_{2,i}) + \dfrac{1-c}{N-1}\sum_{j \neq i} M(\mathbf{z}_j) z_{2,j}}, \tag{A18a}$$

and the probability that individual $i$ sires an ovule of individual $j \neq i$ is given by the share of pollen from

$i$ out of all the pollen received by $j$,

$$\beta(\mathbf{z}_i, \mathbf{z}_j) = \frac{z_{2,i} \frac{1-c}{N-1} M(\mathbf{z}_i)}{M(\mathbf{z}_j)(1-z_{2,j}) + \frac{1-c}{N-1} \sum_{k \neq j} z_{2,k} M(\mathbf{z}_k)}. \tag{A18b}$$

The rest of the life cycle proceeds as explained in Section A.1.1. Self-fertilised zygotes suffer from inbreeding depression, becoming viable with probability $1-\delta$, and viable diploid zygotes immediately undergo meiosis to give rise to two haploid seeds each. Then, all adults die and the next generation is sampled from the produced seeds.

##### A.2.1 Directional selection

Using eq. (A2), the selection gradient on pollen export $z_2$, is given by

$$S_2(\mathbf{z}) = K(\mathbf{z}) \left( \left. \frac{\partial B(\mathbf{z}_\bullet, \mathbf{z})}{\partial z_{2\bullet}} \right|_\circ + (1-2\delta) \left. \frac{\partial \alpha(\mathbf{z}_\bullet)}{\partial z_{2\bullet}} \right|_\circ \right) = \underbrace{K(\mathbf{z}) \frac{1-c}{(1-cz_2)^2}}_{>0} \Big( 1 - z_2(1+c-2\delta) \Big), \tag{A19a}$$

which is independent of sex allocation, as pointed out by Zhang Zhang (2000). This is because the selection gradient captures directional selection on each trait holding everything else fixed, in which case all individuals have the same pollen production $M(\mathbf{z})$, which simplifies away in eq. (A18).

Pollen export $z_2$ does not affect ovule and pollen production ($\left. \frac{\partial F(\mathbf{z}_\bullet)}{\partial z_{2\bullet}} \right|_\circ = \left. \frac{\partial M(\mathbf{z}_\bullet)}{\partial z_{2\bullet}} \right|_\circ = 0$). The selection gradient in eq. (A19a) therefore reflects a balance between the fitness consequences of a change in pollen export through its effect on siring success as a pollen donor and the selfing rate, with the direction of selection on pollen export dictated by the term in brackets on the right-hand side of eq. (A19a). This term is linear in $z_2$, and upon closer inspection reveals that one of two things can happen depending on the cost of dispersal $c$ and inbreeding depression $\delta$. Either inbreeding depression is strong relative to the cost of dispersal, specifically,

$$\delta \geqslant \frac{c}{2}, \tag{A19b}$$

in which case eq. (A19a) is positive for all $z_2 \in [0, 1]$, so that selection favours complete pollen dispersal

($z_2 \to 1$), leading the population to become completely outcrossing; or inbreeding depression is weak relative to the costs of dispersal ($\delta < c/2$), in which case there exists an intermediate pollen export rate

$$z_2^* = \frac{1}{1 + c - 2\delta} \tag{A19c}$$

at which directional selection vanishes. The term between brackets is then strictly positive at $z_2 = 0$ and negative at $z_2 = 1$. Given it is linear in $z_2$, this implies that the singular pollen export rate in eq. (A19c) is always convergence stable. The singular export rate is a decreasing function of the cost of dispersal: the more pollen gets lost during export, the lower selection for pollen dispersal because its benefits in terms of siring success as pollen donor are reduced. Eq. (A19c) is also an increasing function of inbreeding depression: the higher inbreeding depression, the lower the benefits of selfing are, which promotes higher dispersal.

The selection gradient on sex allocation $z_1$, meanwhile, is given by

$$S_1(\mathbf{z}) = K(\mathbf{z}) \Bigg[ \underbrace{\Big(1 + (1 - 2\delta)\alpha(\mathbf{z})\Big)\frac{1}{z_1}}_{\text{Effect on seed production}} - \underbrace{(1 - 2\delta)\alpha(\mathbf{z})\big(1 - \alpha(\mathbf{z})\big)\frac{1}{1 - z_1}}_{\text{Effect on selfing rate}} - \underbrace{\big(1 - \alpha(\mathbf{z})\big)\frac{1}{1 - z_1}}_{\substack{\text{Effect on success} \\ \text{as pollen donor}}} \Bigg] \tag{A20a}$$

where

$$\alpha(\mathbf{z}) = \frac{1 - z_2}{1 - cz_2} \tag{A20b}$$

is the selfing rate in the resident population, which does not depend on sex allocation because all individuals produce the same amount of pollen so that $M(\mathbf{z})$ cancels out in eq. (A18a).

Equation (A20a) shows that directional selection on sex allocation is shaped through the three pathways described in Section A.1.2, which all depend on the selfing rate $\alpha(\mathbf{z})$. The first term, which gives the fitness consequences of a change in sex allocation through its effect on seed production, is always positive (increasing allocation to female function always increases fitness returns through this function). Its influence on sex allocation evolution is amplified by selfing when inbreeding depression is weak ($\delta < 1/2$): it selects more strongly for increased allocation to female function, because selfing increases the fitness returns per seed in this case. Conversely, selfing tends to reduce selection for allocation to female function through this effect under strong inbreeding depression. The second term gives the fitness consequences of a change in sex allocation through its effect on the selfing rate, which depend on the magnitude of inbreeding depression as well. This term is negative when selfing is beneficial ($\delta < 1/2$): it promotes greater allocation to male function because this increases the selfing rate; and positive when selfing is detrimental ($\delta > 1/2$), promoting greater allocation to female function because this reduces the selfing rate. Finally, the third term gives the fitness effect of a change in sex allocation through its effect on success as a pollen donor. It is always negative, therefore promoting allocation to male function (producing more pollen always has a positive impact on fitness returns as a pollen donor), but is always reduced by selfing. This is because the more individuals retain pollen for selfing, the less likely it is that an outcrossing pollen donor will be able to fertilise them.

Re-arranging eq. (A20a), we obtain

$$S_1(\mathbf{z}) = K(\mathbf{z})\Big(1 + (1-2\delta)\alpha\left(\mathbf{z}\right)\Big)\left[\frac{1}{z_1} - \big(1-\alpha\left(\mathbf{z}\right)\big)\frac{1}{1-z_1}\right], \tag{A20c}$$

which tells us that the opposing effects of selfing on fitness returns through a focal individual's own ovules (the first two terms in eq. A20a) exactly balance out: any increase in ovule production is met with a proportional decrease in the expected fitness return per ovule. Thus, only the effect of selfing on success as a pollen donor remains, which tends to favour female-biased sex allocations. Solving for the singular sex allocation (eq. (A3), we find

$$S_1(\mathbf{z}^*) = 0 \quad \Leftrightarrow \quad z_1^* = \frac{1}{2-\alpha\left(\mathbf{z}^*\right)}, \tag{A20d}$$

in agreement with previous analyses (e.g., eq. 31 with $n = 1$ in Zhang (2000); also derived in Lesaffre et al. (2025)).

Combining this result with our findings on pollen export rate (eq. A19), we have that directional selection either favours complete outcrossing ($z_2 \to 1$ when $\delta \geqslant c/2$), in which case it also favours equal investment to male and female function ($z_1^* = 1/2$); or it favours an intermediate export rate $z_2^* = 1/(1 + c - 2\delta)$, resulting in mixed mating and a female-biased sex allocation.

##### A.2.2 Disruptive selection

When inbreeding depression is sufficiently strong for selection to favour complete outcrossing, pollen export is under persistent directional selection to remain complete ($S_2(\mathbf{z}) > 0$ at $z_2 = 1$), so that the model collapses to a randomly mating hermaphroditic population with linear fitness returns through each sexual function. In this situation, equal investment into male and female function, $z_1^* = 1/2$, constitutes an evolutionary stable strategy (Charnov, 1982; Lesaffre et al., 2024, 2025). We therefore do not consider this case further.

In contrast, when inbreeding depression is such that an intermediate rate of pollen export is favoured, directional selection vanishes on both traits at the singular point $\mathbf{z}^* = (z_1^*, z_2^*)$ so that second-order forces as described by the Hessian matrix at $\mathbf{z}^*$ dictate evolutionary dynamics (Section A.1.2.3 above).

Using eq. (A9b), the quadratic selection coefficient on sex allocation, $H_{11}(\mathbf{z}^*)$, is given by

$$H_{11}(\mathbf{z}^*) = \underbrace{K(\mathbf{z}^*)(1-2\delta)}_{\text{Positive when } \delta<c/2} \left[ \left.\frac{\partial^2 \alpha\left(\mathbf{z}_\bullet\right)}{\partial {z_{1\bullet}}^2}\right|_* + 2 \left.\frac{\partial \alpha\left(\mathbf{z}_\bullet\right)}{\partial z_{1\bullet}}\right|_* \times \frac{1}{F(\mathbf{z}^*)} \left.\frac{\partial F(\mathbf{z}_\bullet)}{\partial z_{1\bullet}}\right|_* \right]. \tag{A21a}$$

The term between square brackets depends on two elements. The first is the second derivative of the selfing rate with respect to mutant sex allocation. This is strictly negative, because the individual's own pollen increasingly competes with itself for self-fertilisation as the amount produced increases, leading to diminishing returns on the selfing rate (analogous to a form of local sperm/mate competition, Hamilton, 1967; Schärer, 2009). The second term captures the joint effect of a change in sex allocation on ovule production and the selfing rate. Since any change in sex allocation has opposite effects on these two quantities, this term is necessarily negative as well. Thus, eq. (A21a) as a whole is negative, indicating that partial selfing tends to generate stabilising selection on sex allocation under weak inbreeding depression (Lesaffre et al., 2025).

The quadratic selection coefficient on pollen export, $H_{22}(\mathbf{z}^*)$, is given by

$$H_{22}(\mathbf{z}^*) = \underbrace{K(\mathbf{z}^*)(1-2\delta)}_{>0} \left. \frac{\partial^2 \alpha\left(\mathbf{z}_\bullet\right)}{\partial {z_2}_\bullet^{\,2}} \right|_*, \tag{A21b}$$

so that its sign is determined by the sign of $\left. \frac{\partial^2 \alpha(\mathbf{z}_\bullet)}{\partial {z_2}_\bullet^{\,2}} \right|_*$, which is also negative for reasons similar to those exposed above: as the proportion of pollen exported decreases, competition amongst the individual's own pollen grains intensifies and leads to diminishing returns in terms of increased selfing. Quadratic selection on pollen export is therefore also stabilising.

Finally, the correlational selection coefficient between sex allocation and pollen export, $H_{12}(\mathbf{z}^*)$, is

$$H_{12}(\mathbf{z}^*) = \underbrace{K(\mathbf{z}^*)}_{>0} \left[ \left. \frac{\partial^2 B(\mathbf{z}_\bullet, \mathbf{z})}{\partial {z_1}_\bullet \partial {z_2}_\bullet} \right|_* + (1-2\delta) \left( \left. \frac{\partial^2 \alpha\left(\mathbf{z}_\bullet\right)}{\partial {z_1}_\bullet \partial {z_2}_\bullet} \right|_* + \left. \frac{\partial \alpha\left(\mathbf{z}_\bullet\right)}{\partial {z_2}_\bullet} \right|_* \times \frac{1}{F(\mathbf{z}^*)} \left. \frac{\partial F(\mathbf{z}_\bullet)}{\partial {z_1}_\bullet} \right|_* \right) \right]. \tag{A21c}$$

The first term in square brackets captures the effect of the interaction between the two traits on success as a pollen donor, which is always negative. This is because increasing allocation to female function $z_1$ always has a negative impact on success as a pollen donor, and this impact becomes increasingly strong (i.e., more negative) as the focal individual's pollen export increases. In other words, a higher pollen export rate makes a decrease in pollen production more negative for siring success as a pollen donor. This term therefore tends to favour a negative correlation between sex allocation and pollen export.

The second term is proportional to $1 - 2\delta$, capturing the effect of the interaction between $z_1$ and $z_2$ on fitness gains through the focal individual's own ovules, which is decomposed in two parts. The first part is the cross-derivative of the selfing rate with respect to both traits $\left. \frac{\partial^2 \alpha(\mathbf{z}_\bullet)}{\partial {z_1}_\bullet \partial {z_2}_\bullet} \right|_*$, which captures the effect of the interaction between the two traits on the selfing rate of a focal individual. This term is positive when the singular pollen export rate is above a threshold, specifically such that

$$\alpha\left(\mathbf{z}^*\right) < \frac{1}{2} \quad \Leftrightarrow \quad c < \frac{1+2\delta}{2}, \tag{A21d}$$

which promotes a positive association between sex allocation and pollen export. This is because in this case, the pollen retained by the individual makes up the minority of the total amount of pollen that it receives, so that increasing pollen export has a small impact on this total amount. Increasing the pollen export rate then tends to decrease the negative impact that increasing the focal individual's allocation to female function has on its own selfing rate, which favours a positive correlation between the two traits (the contribution of the individual's own pollen to the denominator in eq. A18a is relatively small, so that the effect of traits changes is mostly driven by the numerator). Conversely, this term promotes a negative association between sex allocation and pollen export when the pollen export rate is below the threshold. This is because an individual's own pollen then makes up the majority of the total amount of pollen it receives, so that increasing its pollen export rate decreases both the amount of pollen it retains and the total amount of pollen received, which overall tends to amplify the effect of the focal's sex allocation on its selfing rate. The second part captures the joint effect of a change in pollen export on the selfing rate ($\frac{\partial \alpha(\mathbf{z}_\bullet)}{\partial z_{2\bullet}}\Big|_\circ$) and a change in sex allocation on relative ovule production ($\frac{\partial F(\mathbf{z}_\bullet)}{\partial z_{1\bullet}}\Big|_\circ / F(\mathbf{z}^*)$) because increasing pollen export has a negative effect on the selfing rate and increasing sex allocation has a positive effect on ovule production, this part is negative and therefore promotes a negative correlation between pollen export and sex allocation. Overall, plugging eq. (A18) into eq. (A21c), the correlation selection coefficient simplifies to

$$H_{12}(\mathbf{z}^*) = -\frac{(1-c)(1+c-2\delta)}{2z_1^*(1-z_1^*)\left[1-\delta\big(c+2(1-\delta)\big)\right]}\left(1+\frac{1-2(c-\delta)}{c-2(1-\delta)}\right), \tag{A21e}$$

which is strictly negative for $\delta < c/2$, indicating that correlational selection on sex allocation and pollen export overall favours a negative correlation between the two traits, i.e., it favours individuals that allocate more to their male function to export pollen at a higher rate, and vice-versa.

Putting the elements of the Hessian matrix together (eq. A21), we compute the characteristic polynomial $\mathbf{H}(\mathbf{z}^*)$, the roots of which give the eigenvalues of this matrix and therefore inform us on the nature of selection at $\mathbf{z}^*$. We refrain from giving its full expression here as it is unsightly, but it is available in the accompanying *Mathematica* notebook (`AnalyticalResults.nb` available at `10.5281/zenodo.23062245`). The polynomial takes the form

$$\mathcal{P}(\lambda) = \lambda^2 + \gamma_1\lambda + \gamma_0, \tag{A22}$$

and is therefore an upward-facing parabola, as its quadratic coefficient is positive. This implies that the parabola is negative between its two roots. Thus, if it is negative at $\lambda = 0$, then exactly one eigenvalue of $\mathbf{H}(\mathbf{z}^*)$ is positive. Evaluating $\mathcal{P}(\lambda)$ at this point we have

$$\mathcal{P}(0) = \gamma_0 = -\left[\frac{(1-c)(1+c-2\delta)\big(c-2(1-\delta)\big)}{2(1-2\delta)\left[1-\delta\big(c+2(1-\delta)\big)\right]}\right]^2 < 0. \tag{A23}$$

The Hessian matrix $\mathbf{H}(\mathbf{z}^*)$ therefore always has a positive eigenvalue despite having a strictly negative diagonal when $\delta < c/2$ (Fig. 1A). Biologically, this implies that selection is always disruptive on sex allocation and pollen export as a result of correlational selection, favouring the emergence of two types of individuals: one that allocates more resources to its female function while reducing its pollen export rate, and one that allocates more to its male function with an increased pollen export (Fig. 1B). In other words, selection favours the emergence of one type that specialises in the production of ovules and self-fertilisation, and another that specialises into pollen production and export.

##### A.2.3 Polymorphism resulting from evolutionary branching

We ran numerical analyses as described in Section A.1.3.2 to investigate the nature of polymorphism maintained at evolutionary equilibrium. We find that evolutionary branching results in the coexistence of two types, a pure male type that exports all its pollen ($z_1^* = 0$ and $z_2^* = 1$) and a hermaphroditic type that exports none of its pollen ($z_2^* = 0$), leading to a form of androdioecy. See main text for more details.

#### A.3 Joint evolution with self-pollen advantage

In this section, we derive the analytical results presented in Section 4 of the main text. We investigate the joint evolution of three traits: sex allocation $z_1$ and the pollen export rate $z_2$, as in the previous section, and the self-pollen advantage, $z_3 \in [0, 1]$ which controls the competitive advantage that a plant grants its own pollen relative to pollen from other donors. The population follows the life cycle described in Section A.1.1, and the three traits affect the behaviour of individuals along this life cycle as follows.

*(i) Sexual development:* The number of ovules and pollen grains produced by individual $i$ are given by

$$F(\mathbf{z}_\mathrm{i}) = F_0 z_{1,i} \quad \text{and} \quad M(\mathbf{z}_\mathrm{i}) = M_0(1 - z_{1,i}), \tag{A24a}$$

respectively. We assume that the number of ovules is large relative to the number of breeding spots ($N \ll F_0$), and that there is always enough pollen for all ovules to be fertilised ($F_0 \ll M_0$), i.e., no pollen limitation.

*(ii) Mating:* After sexual development, individuals export a fraction $z_2 \in [0, 1]$ of their pollen to fertilise ovules on other plants, and the rest $(1 - z_2)$ remains on the plant for self-fertilisation (Holsinger, 1991; Zhang, 2000; Porcher and Lande, 2005). A fraction $c \in [0, 1]$ of the exported pollen is lost during dispersal, and the successful fraction $1 - c$ is distributed equally among the $N - 1$ other plants. The self- and cross-pollen received by each plant then compete for ovule fertilisation, and this competition is biased by the maternal plant either in favour of or against its own pollen according to its self-pollen advantage $z_{3,i}$. Competition is unbiased when $z_{3,i} = 1/2$, so that pollen grains from all sources are equally competitive, whereas $z_{3,i} > 1/2$ and $z_{3,i} < 1/2$ favour and oppose self-fertilisation, respectively. The self-pollen advantage $z_3$ thus modulates the relative competitiveness of self- and cross-pollen grains for fertilisation. Overall, the selfing rate $\alpha(\mathbf{z}_i)$ of individual $i$ is given by

$$\alpha(\mathbf{z}_i) = \frac{z_{3,i} M_\mathrm{self}(\mathbf{z}_i)}{z_{3,i} M_\mathrm{self}(\mathbf{z}_i) + (1 - z_{3,i}) \times \underbrace{\frac{1-c}{N-1} \sum_{j \neq i} M_\mathrm{out}(\mathbf{z}_j)}_{\substack{\text{Total amount of cross-pollen} \\ \text{received by plant } i}}}, \tag{A24b}$$

and the probability that individual $i$ fertilises an ovule of individual $j \neq i$ is

$$\beta(\mathbf{z}_i, \mathbf{z}_j) = \frac{(1 - z_{3,j}) \dfrac{1-c}{N-1} M_\mathrm{out}(\mathbf{z}_i)}{z_{3,j} M_\mathrm{self}(\mathbf{z}_j) + (1 - z_{3,j}) \dfrac{1-c}{N-1} \sum_{k \neq j} M_\mathrm{out}(\mathbf{z}_k)}, \tag{A24c}$$

where

$$M_\mathrm{self}(\mathbf{z}_i) = M(z_{1,i})(1 - z_{2,i}) P_\mathrm{self} \quad \text{and} \quad M_\mathrm{out}(\mathbf{z}_i) = M(z_{1,i}) z_{2,i} P_\mathrm{out}(z_{3,i}) \tag{A24d}$$

respectively give the amount of self-pollen transferred to plant $i$'s own stigmas and the amount of pollen exported away by plant $i$. In eq. (A24d), $P_{\text{self}} \in [0,1]$ is a parameter that captures the efficiency of within-plant pollen transfers, while $P_{\text{out}}(z_{3,i}) \in [0,1]$ is function that captures the efficiency of pollen export, which decreases with the individual's self-pollen advantage $z_{3,i}$, reflecting sexual interference and pollen discounting.

Since $M_{\text{self}}(\mathbf{z}_i)$ and $M_{\text{out}}(\mathbf{z}_i)$ appear exclusively in ratios in eqs. (A24b) and (A24c), only their relative scaling matters. Without loss of generality, we may therefore divide both by $P_{\text{self}}$ so that they become

$$M_{\text{self}}(\mathbf{z}_i) = M(z_{1,i})(1 - z_{2,i}) \quad \text{and} \quad M_{\text{out}}(\mathbf{z}_i) = M(z_{1,i}) z_{2,i} P(z_{3,i}), \tag{A24e}$$

where $P(z_{3,i}) = P_{\text{out}}(z_{3,i})/P_{\text{self}}$ is the relative efficiency of pollen export, which we will work with from now on.

The rest of the life cycle proceeds as explained in Section A.1.1. Self-fertilised zygotes suffer from inbreeding depression, becoming viable with probability $1 - \delta$, and viable diploid zygotes immediately undergo meiosis to give rise to two haploid seeds each. Then, all adults die and the next generation is sampled from the produced seeds.

##### A.3.1 Cost-free self-pollen promotion ($P(z_3) = 1$)

We first consider the case where self-pollen promotion causes no sexual interference, so that $P(z_3) = 1$ for all $\mathbf{z}$. In this case, we find that the selection gradient on self-pollen advantage is given by

$$S_3(\mathbf{z}) = K(\mathbf{z}) \frac{z_2(1-c)(1-z_2)}{\left[z_2(1-c)(1-z_3) + (1-z_2)z_3\right]^2}(1 - 2\delta), \tag{A25a}$$

which does not depend on sex allocation $z_1$ for reasons similar to the pollen export rate above (see eq. A19 for explanation). Eq. (A25a) shows that the self-pollen advantage is only under selection when the pollen export rate is intermediate (i.e., when $0 < z_2 < 1$). This is because self-pollen never competes with cross-pollen otherwise, so that $z_3$ is effectively neutral here. Provided that $0 < z_2 < 1$, eq. (A25a) also shows that there can be no intermediate equilibrium for the self-pollen advantage under directional selection, as the gradient never changes sign for $z_3 \in [0,1]$. Thus, at an equilibrium of directional selection, either $z_3^* = 0$ and the population is strictly outcrossing (self-pollen cannot fertilise) unless no pollen is exchanged ($z_2 = z_3 = 0$) which is a degenerate case where the population cannot persist, as no ovule gets fertilised; or $z_3^* = 1$ and the population is strictly selfing (only self-pollen can fertilise) unless individuals export all their pollen ($z_2 = z_3 = 1$) which is also a degenerate case where no ovule gets fertilised and the population goes extinct. Which of these two equilibria will be reached by an evolving population is determined by inbreeding depression. Specifically, either $\delta < 1/2$ and eq. (A25a) is positive, so that a population with an intermediate initial selfing rate will evolve towards complete selfing ($z_3 \to 1$); or $\delta > 1/2$ and eq. (A25a) is negative, so that the population will evolve complete outcrossing ($z_3 \to 0$). The $\delta = 1/2$ threshold comes from the two-fold transmission advantage of selfing which must be overcome by inbreeding depression for outcrossing to be favoured, as shown by classical theory in the absence of sexual interference (Lande and Schemske, 1985).

Once the population has become either completely outcrossing or completely selfing, the joint evolution of sex allocation and pollen export is trivial. Under enforced outcrossing ($z_3 = 0$), the selection gradient on pollen export reduces to

$$S_2(\mathbf{z}) = \frac{K(\mathbf{z})}{z_2}, \tag{A25b}$$

which is always positive, and so favours complete pollen dispersal. This is because any pollen that is not dispersed is wasted when self-fertilisation cannot occur. The selection gradient on sex allocation, meanwhile, is given by

$$S_1(\mathbf{z}) = K(\mathbf{z}) \left( \frac{1}{z_1} - \frac{1}{1 - z_1} \right), \tag{A25c}$$

which cancels at $z_1^* = 1/2$, as expected in a randomly mating population.

Under enforced selfing ($z_3 = 1$), the selection gradient on pollen export is zero ($S_2(\mathbf{z}) = 0$) because pollen export does not affect reproductive success: pollen dispersal is neutral. The selection gradient on sex allocation is given by

$$S_1(\mathbf{z}) = K(\mathbf{z}) \frac{2(1 - \delta)}{z_1}, \tag{A25d}$$

which is positive, indicating that selection favours allocating all resources to female function. This is because the amount of pollen produced does not matter for reproductive success as any non-zero production is sufficient for self-fertilisation, so that individuals producing more ovules are always at an advantage. Eventually, however, this should lead the population to produce no pollen at all and so to go extinct. This degenerate case stems from our assumption that pollen is never limiting for ovule fertilisation. In a more elaborate model of mating where the number of pollen grains required for fertilisation of available ovules is explicitly accounted for, the population would likely stabilise at a small but non-zero production of pollen, as seen in highly selfing species exhibiting a selfing syndrome.

##### A.3.2 Sexual interference caused by self-pollen promotion

The above shows that mixed mating is never favoured by directional selection when maternal plants can manipulate the outcome of pollen competition without incurring any cost, consistent with existing theory (e.g., Lande and Schemske, 1985; Abu Awad and Roze, 2020). We now investigate the effect of sexual interference on the joint evolution of self-pollen advantage, pollen export and sex allocation.

###### A.3.2.1 Directional selection

Under sexual interference, the selection gradient on self-pollen advantage is given by

$$\begin{aligned} S_3(\mathbf{z}) = & \underbrace{K(\mathbf{z})\frac{z_2(1-c)(1-z_2)P(z_3)}{\left[z_2(1-c)(1-z_3)P(z_3)+(1-z_2)z_3\right]^2}}_{\geqslant 0} \\ & \times\left[1-2\delta+(1-z_3)\left(z_3+\frac{z_2(1-c)}{1-z_2}(1-z_3)P(z_3)\right)\frac{P'(z_3)}{P(z_3)}\right], \end{aligned} \tag{A26}$$

which does not depend on sex allocation, as before, and now features an extra term that depends on the sexual interference function $P(z_3)$ and its first derivative ($P'(z_3)$). This term is negative when self-pollen promotion causes sexual interference (i.e., when $P'(z_3) < 0$) and so opposes selection for self-fertilisation owing to the transmission advantage when inbreeding depression is weak ($\delta < 1/2$), indicating that sexual interference could in principle allow the existence of an intermediate equilibrium self-pollen advantage, potentially leading to mixed mating.

The selection gradient on pollen export, meanwhile, is given by

$$S_2(\mathbf{z}) = \underbrace{K(\mathbf{z})\frac{(1-c)(1-z_3)P(z_3)}{\left[z_2(1-c)(1-z_3)P(z_3)+(1-z_2)z_3\right]^2}}_{\geqslant 0}\left[z_3 - z_2\Big(2(1-\delta)z_3 - P(z_3)(1-c)(1-z_3)\Big)\right]. \tag{A27a}$$

The term on the right-hand side is always positive when $z_3 < 1$ and so plays no role in determining the direction of selection. The term on the second line is linear in $z_2$. When $z_2 = 0$, the gradient reduces to

$$S_2(\mathbf{z}) = K(\mathbf{z})\frac{P(z_3)(1-c)(1-z_3)}{z_3} \tag{A27b}$$

which is always positive for $0 \leqslant z_3 < 1$, indicating that selection always favours some pollen export provided that outcrossing is possible. Therefore, if $S_2(\mathbf{z}) < 0$ when $z_2 = 1$, then there is a single singular pollen export value to which the population will converge under directional selection. Otherwise, selection favours complete pollen export ($z_2 \to 1$). When $z_2 = 1$, the gradient becomes

$$S_2(\mathbf{z}) = K(\mathbf{z})\left[1 - \frac{z_3(1-2\delta)}{(1-c)(1-z_3)P(z_3)}\right], \tag{A27c}$$

which is always positive when $\delta \geqslant 1/2$. Selection thus favours the evolution of complete pollen export ($z_2 \to 1$) when $\delta \geqslant 1/2$, that is, when inbreeding depression outweighs the transmission advantage of selfing. Otherwise, whether the gradient turns negative depends on the value of $z_3$.

One can show that because $P'(z_3) < 0$, eq. (A27c) is a strictly decreasing function of $z_3$ for $\delta < 1/2$, is strictly positive when $z_3 = 0$ and tends to $-\infty$ as $z_3$ approaches 1, i.e.,

$$\lim_{z_3 \to 1^-} S_2(\mathbf{z}) = -\infty. \tag{A27d}$$

These properties imply that the gradient on pollen export at $z_2 = 1$ is negative above a critical value $z_3' \in [0,1)$. Thus, when $z_3' < z_3 < 1$ there is a singular intermediate pollen export rate $z_2^* \in (0,1)$ and otherwise selection favours complete export ($z_2 \to 1$). When $z_3 > z_3'$, the singular pollen export rate is given by

$$z_2^* = \frac{z_3}{2(1-\delta)z_3 - P(z_3)(1-c)(1-z_3)}. \tag{A27e}$$

Overall, The analysis presented in (A27) shows that either complete pollen export evolves, or selection favours an intermediate pollen export rate (eq. A27e), which occurs when $\delta < 1/2$ and $z_3' < z_3 < 1$ (Note that when $z_3 = 1$, the pollen export rate is effectively neutral as only self-pollen can fertilise ovules).

**Complete outcrossing is always stable.** When $z_2 \to 1$, the selection gradient on self-pollen advantage (eq. A26) becomes

$$S_3(\mathbf{z}) = K(\mathbf{z})\frac{P'(z_3)}{P(z_3)}, \tag{A28}$$

which is strictly negative (since $P'(z_3) < 0$). This shows that in the presence of sexual interference, a population that is initially exporting pollen at a high rate will be driven towards a self-pollen advantage of zero. In the vicinity of $z_2 = 1$ and $z_3 = 0$, we thus have that $S_2(\mathbf{z}) = 1 > 0$ and $S_3(\mathbf{z}) < 0$, which shows that selection always maintains the population at this fully outcrossing strategy, making it an absorbing state of gradual evolutionary dynamics at which evolution comes to a halt, because directional selection does not vanish on either trait. The reason this holds even when inbreeding depression is absent is that the two traits interfere with one another. When $z_3$ is close to zero, reducing $z_2$ to retain pollen for self-fertilisation leads to such a small increase in selfing rate that this benefit is always trumped by the cost incurred in terms of lost mating success as pollen donor. When $z_2$ is close to one, meanwhile, increasing $z_3$ leads to a loss of pollen export under sexual interference that always outweighs the potential benefits in terms of increased selfing. Thus, enforced outcrossing is an evolutionary trap that cannot be escaped under sexual interference.

**Complete selfing is also stable.** When $z_3 = 1$, so that only self-pollen can fertilise maternal plants, the gradient on self-pollen advantage reduces to

$$S_3(\mathbf{z}) = K(\mathbf{z})\frac{z_2(1-c)}{1-z_2}(1-2\delta)P(1), \tag{A29}$$

which is strictly positive when $\delta < 1/2$, showing that $z_3$ is always selected to increase when it is sufficiently close to one under weak inbreeding depression. In the vicinity of $z_3 = 1$, the pollen export rate is also driven towards an intermediate $z_2^*$ (eq. A27e). Thus, a population with an initially high self-pollen advantage will be selected to remain so, showing that complete selfing is also a stable equilibrium of directional selection under sexual interference.

**Mixed mating can be favoured.** One or more stable mixed mating equilibria could exist between the complete selfing and complete outcrossing equilibria, depending on the shape of the sexual interference function $P(z_3)$. However, explicitly showing the existence of such intermediate stable points becomes difficult even in the simplest of cases because eq.(A26) depends on $z_3$ in a complicated way. We therefore turn to numerical analyses, which requires picking a shape for $P(z_3)$. For the rest of our analyses, we assume that $P(z_3)$ is given by

$$P(z_3) = \frac{2\left(1 - e^{\kappa/2}\right)}{2 - e^{\kappa/2} - e^{\kappa z_3}}. \tag{A30}$$

Equation (A30) is a decreasing sigmoid function that satisfies $P(1/2) = 1$, so that our previous model without maternal control on pollen competition is recovered for $z_3 = 1/2$. The shape of $P(z_3)$ is controlled by parameter $\kappa > 0$. The higher $\kappa$, the more intense sexual interference is caused by self-pollen promotion.

We analysed the outcome of directional selection numerically using the approach described in section A.1.2.2, results are presented in Figure 2B-D in the main text.

###### A.3.2.2 Disruptive selection

The complete selfing and complete outcrossing equilibria are endpoints of gradual evolution that need not be given further consideration. In contrast, the mixed mating equilibrium (where it exists) is a singular point at which second-order forces may act. We computed the leading eigenvalue of the Hessian matrix $\mathbf{H}(\mathbf{z}^*)$ at this equilibrium and found that it is never evolutionarily stable, as the leading eigenvalue of $\mathbf{H}(\mathbf{z}^*)$ is always positive. See Main Text Section 4.3 for details.

#### Appendix B

### Individual-based simulations

In this section, we describe the individual-based simulation program that we used to obtain the results presented in Figs. 1B and 3B-C of the main text. The program is coded in C++20 and is available at: `10.5281/zenodo.23062245`.

It depicts a population of $N$ haploid individuals characterised by their allelic values for three traits: sex allocation $z_1$, pollen export rate $z_2$ and self-pollen advantage $z_3$. Each generation proceeds as follows.

**Fecundities and selfing rates.** We first determine the trait values expressed by each individual by clipping their allelic values so that they lie between 0 and 1. From these, we compute the female fecundity of individuals as

$$F(z_{1,i}) = z_{1,i} \tag{B1}$$

for individual $i \in \{1, \ldots, N\}$; their contribution to the outcross pollen pool, given by

$$M_{\text{out}}(\mathbf{z}_i) = (1 - z_{1,i}) z_{2,i} P(z_{3,i}), \tag{B2}$$

where

$$P(z_{3,i}) = \frac{2(1 - e^{\kappa/2})}{2 - e^{\kappa/2} - e^{\kappa z_{3,i}}} \tag{B3}$$

denotes pollen discounting caused by the self-pollen advantage, and their selfing rate,

$$\alpha(\mathbf{z}_i) = \frac{z_{3,i} M_{\text{self}}(\mathbf{z}_\text{i})}{z_{3,i} M_{\text{self}}(\mathbf{z}_\text{i}) + (1 - z_{3,i}) \dfrac{1-c}{N-1} \sum_{j \neq i} M_{\text{out}}(\mathbf{z}_\text{j})}, \tag{B4}$$

where $M_{\text{self}}(\mathbf{z}_i) = (1 - z_{1,i})(1 - z_{2,i})$.

**Offspring production.** To create the next generation, we create $N$ offspring as follows. For each offspring, we first pick a mother with replacement from the current generation. Each individual $i$ has a probability $F(z_{1,i})/\sum_j F(z_{1,j})$. We determine whether the produced offspring is selfed (with probability $\alpha(\mathbf{z}_i)$) or outcrossed (with probability $1 - \alpha(\mathbf{z}_i)$). If it is selfed, we mark it as such and assign it a viability $w = w_{\text{self}} = 1 - \delta$. If it is outcrossed, we also mark it and assign it a viability $w = w_{\text{out}} = 1$. We then check whether the produced offspring survives (with probability $w$). If it does not survive, we discard it and start over by sampling a new mother, determining whether the new offspring is selfed and so on. We repeat this process until a surviving offspring is obtained.

**Haplotype transmission.** If this surviving offspring is selfed, then it inherits its mother's haplotype. If it is outcrossed, it either inherits the mother's haplotype (with probability $1/2$) or that of its father, in which case said father must be sampled from the population. To do so, we pick individuals from the current generation, with the probability that individual $i$ is chosen given by $M_{\text{out}}(\mathbf{z}_i)/\sum_j M_{\text{out}}(\mathbf{z}_j)$. If the chosen individual turns out to be the same as the mother, it is discarded and a new father is picked until we find one that is different from the maternal individual.

**Mutation.** Once the offspring has been constructed, it undergoes mutation with probability $\mu$, in which case all the allelic values are changed by a small deviation sampled independently in a normal distribution with mean zero and standard deviation $\sigma$. Allelic values are clamped to remain between $-0.2$ and $1.2$. Allowing allelic values to vacate the $[0, 1]$ interval over which traits are defined prevents the occurrence of directional mutational bias due to boundary effects.

**Measurements.** Every $t_{\text{mes}}$ generations, we record the allelic values for all traits and the selfing rates of $n_{\text{mes}}$ randomly sampled individuals in the population.

**Initialisation.** The population is initially fixed for phenotype $\mathbf{z}_{\text{init}} = (z_{1,\text{init}}, z_{2,\text{init}}, z_{3,\text{init}})$.
